# Autoinduction expression systems via engineered quorum-sensing circuits in *Synechococcus elongatus PCC 7942*

**DOI:** 10.1101/2025.04.03.647055

**Authors:** Emmanuel J. Kokarakis, María Santos-Merino, Sajjad Ghaffarinasab, Daniel Vocelle, Daniel C. Ducat

## Abstract

Despite significant potential for cyanobacteria as sustainable bioproduction chases, there are limited examples of scaled cyanobacterial bioproduction. In part, this is because most cyanobacterial species are poorly adapted to bioreactor cultivation conditions and lack features that facilitate biomass growth and harvesting at scale. We explored quorum sensing (QS) pathways derived from heterotrophic microbes as a method for autoinduction of gene expression circuits coordinated to population density in cyanobacteria. Here, we integrated genetic modules designed to produce and detect the diffusible QS signal, acyl-homoserine lactones (AHLs), in the cyanobacterial model, *Synechococcus elongatus* PCC 7942 (*S. elongatus*). We demonstrate that *S. elongatus* heterologously produces sufficient AHL signals to activate gene expression in a dose-dependent and population density-responsive manner. A hybrid combination of AHL synthesis enzyme from *Vibrio fischeri* (Lux system) with the transcription factor receiver from *Pseudomonas aeruginosa* (Las system) provides an ideal activation ratio and mitigates toxicity observed with some AHL systems. As a proof of concept, we coupled the QS pathway to the expression of a cell division inhibitory gene, *cdv3*, facilitating late-phase cell elongation, cell sedimentation, and improved biomass recovery. Our findings provide a foundation for the development of auto-induction systems leverageable to improve cyanobacterial biotechnology applications.

## 1. Introduction

Quorum sensing (QS) is a broadly conserved mechanism bacteria utilize to coordinate population-level behaviors. QS relies on the production, release, and subsequent detection of signaling molecules known as autoinducers. In turn, autoinducers are recognized by specific receiver proteins that bind them and activate or suppress gene transcription^1,2^. In this manner, increasing cell density leads to accumulation of autoinducers in the surrounding environment until they reach a threshold level that triggers a synchronized, population-wide response through the induction of gene expression. QS pathways are well-characterized in many heterotrophic bacteria for their governance of coordinated cell behaviors, including bioluminescence, biofilm formation, virulence factor production, and other lifestyle transitions^3–6^. QS behaviors are broadly observed across many bacteria, although the specific chemical composition of autoinducers and QS receptor proteins vary across species^1,7,8^.

Among the QS systems, the most extensively studied are derived from Gram-negative bacteria and involve autoinducer molecules known as acyl homoserine lactones (AHLs). AHLs are characterized by a lactone ring linked to a side chain comprising 4 to 18 acyl carbons^9^. The small and apolar characteristics of AHLs allow them to permeate the bacterial cell membrane without a dedicated transporter or transmembrane receptor^4,10,11^. Originally characterized in *Vibrio fischeri*, where the QS system regulates the activation of the luciferase operon *luxICDABE* and bioluminescence^12^, the LuxI-family of proteins catalyze the production of AHLs that control diverse QS behaviors across prokaryotes^1,11,13^. The LuxR family cytosolic proteins detect AHL levels via a mechanism involving AHL binding within a hydrophobic pocket located at the N-terminus, which stabilizes the tertiary structure of this domain, promoting dimerization and recruitment of the transcription factor to gene promoters^14–16^.

The autoinduction and density-dependent properties natively found in QS systems are useful for many biotechnology applications^17–19^. For example, QS systems have been used to study population dynamics, synchronize cell lysis for drug delivery and coordinate late activation of genes to improve the target molecular bioproduction^20–22^. However, a significant knowledge gap remains regarding cyanobacterial QS mechanisms. While some cyanobacteria exhibit population-level behaviors that appear phenotypically analogous to known QS pathways, most cyanobacteria do not encode complete genetic circuits of known signaling pathways such as the AHL systems^23,24^. While mixed communities containing cyanobacteria contain QS autoinducer molecules, there are only limited studies implicating pathways such as biofilm formation and microcystin production to QS-like signaling^25–27^. For example, accumulation of AHLs is associated with biofilm formation in *Microcystis aeruginosa*^28,29^ and *Gloeothece* sp. PCC 6909 produces 3OC8-HSL that may be tied to expression of ribulose-1,5-bisphosphate carboxylase/oxygenase (Rubisco)^23^. However, cyanobacterial QS examples are few and homologs of well-studied QS pathways are not widely encoded in most species^30^.

Installation of heterologous QS systems in cyanobacteria could be useful for self-activation of engineered genetic pathways which would otherwise be cost-prohibitive to control via addition of chemical inducers. Additionally, some problems of cyanobacterial cultivation are relatively universal and apply across most species and irrespective of the target products/services to be produced in large-scale applications. For example, efficient harvesting of cyanobacterial biomass and dewatering costs can account for ∼30% of total cultivation costs, significantly impacting economic competitiveness for cyanobacterial derived bioproducts^31,32^. Additionally, cyanobacteria have recalcitrant cell walls that require harsh solvents or mechanical stress to lyse, which also increases costs of bioproduct extraction^33,34^. Genetic circuits could address these types of challenges, but engaging engineered pathways at the right time paramount, and currently available chemical inducers are expensive, consisting of up to 60% of input costs for growth media^35^. In contrast, QS pathways hold the potential to allow for autoinduction of genetic pathways at defined growth stages of the culture.

Here, we explore the utilization of QS circuits in the model cyanobacterium, *S. elongatus* PCC 7942 (*S. elongatus*), and evaluate their potential to trigger engineered genetic pathways at specific biomass stages. We characterize three different QS systems for favorable features in terms of AHL production rates, cell toxicity, capacity to trigger cognate promoter elements in dose dependent responses, and timing of engineered QS output genes. We show that *S. elongatus* can be programmed to produce sufficient amounts of AHLs to activate QS circuits and confirm that the cell density influences the kinetics of system expression. As proof of concept, we couple QS activation with a cell elongation-inducing genetic pathway with the goal of improving cellular physiology for harvesting, dewatering, and bioproduct recovery. We find *S. elongatus* cell length and sedimentation rates can be greatly increased in late phase cultures using customized QS systems, providing a potential framework for other programmable quorum-like behaviors in cyanobacterial strains.

## 2. Results

### 2.1. *S. elongatus* can synthesize and secrete AHLs

We first interrogated the capacity of *S. elongatus* to produce and secrete acyl-homoserine lactone (AHL) autoinducer molecules. We focused on the production of 3OC6-HSL and 3OC12-HSL, which displayed more favorable properties for an inducible genetic platform in *S. elongatus* based on our recent analysis^36^. We constructed two strains encoding the cognate AHL synthetases LuxI and LasI for 3OC6-HSL and 3OC12-HSL synthesis, respectively^37,38^. Coding sequences for *luxI* and *lasI* were placed under the control of an IPTG-inducible P*_trc_* promoter (Fig. 1a) to allow experimental control of timing and magnitude of gene expression. We followed the appearance of 3OC6-HSL (for *luxI*) and 3OC12-HSL (*lasI*) in the supernatant, specifically detecting the corresponding AHL molecules by liquid chromatography-mass spectrometry (LC-MS; Figs. 1b and 1c).

**Figure 1.**
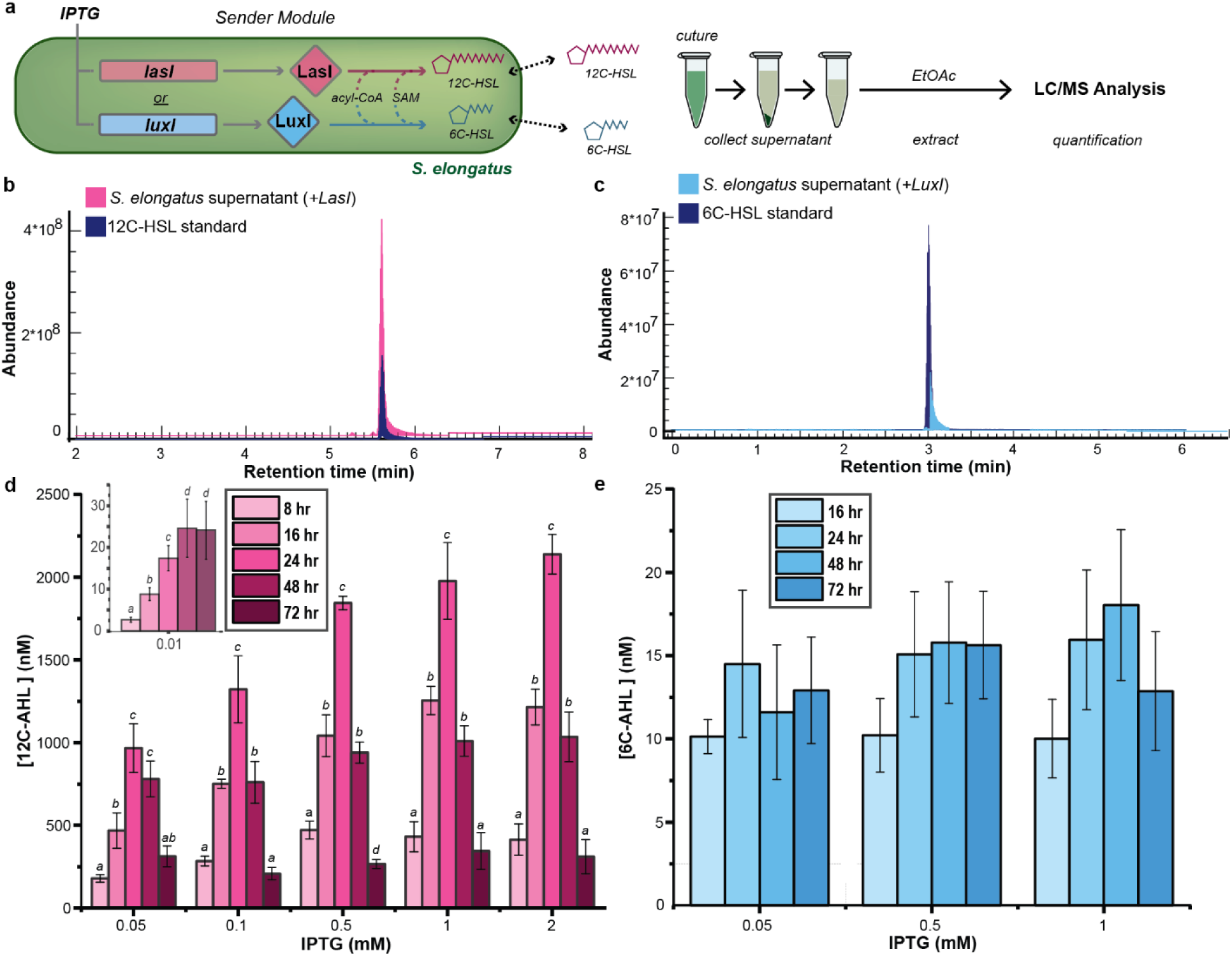
AHL detection and characterization in *S. elongatus*. Overview of diagram showing the strains and the workflow of AHL extraction and detection (a). AHL extracted from *S. elongatus* supernatant and quantified using LC-MS as compared to overlaid chromatograms of AHL standards (b) 3OC12-HSL and (c) 3OC6-HSL. Biosynthesis of 3OC12-HSL (d) and 3OC6-HSL (e), extracted from *Synechococcus* strains expressing LasI and LuxI, respectively. AHLs were extracted from the supernatant overtime after the indicated time following induction across a range of IPTG concentrations 0.01 to 2 mM. Averages of ≥3 independent biological replicates are shown ± SD. Significance was calculated by one-way ANOVA followed by Tukey’s multiple comparison test. Bars labeled with different letters are significantly different (*P* < 0.05).

We evaluated the temporal dynamics of AHL accumulation in the culture medium over 72 hours as a function of varied expression levels of LasI and LuxI (0.005 mM to 2 mM IPTG). Following *lasI* induction, 3OC12-HSL accumulated in the supernatant in a time-dependent and IPTG-dependent manner (Fig. 1d; Supplemental Fig. S1). 3OC12-HSL was detected as early as 8 hours post-induction, and its concentration continued to rise over the first 24 hours (Fig. 1d). Above the minimal threshold [ITPG] of 0.01 mM, *lasI* induction was strongly correlated to 3OC12-HSL production: with peak AHL measurements 24 hours after induction ranging from 1000 nM to 2200 nM. Uninduced cultures did not show any accumulation of AHLs within our detection limits (1 nM). In contrast, strains expressing *luxI* produced 3OC6-HSL at concentrations approximately two orders of magnitude lower and did not show significant trends in AHL production with increasing induction (Fig. 1e). The relatively weak 3OC6-HSL production suggests that there may be other limiting factors in *S. elongatus*, such as substrate availability of short chain fatty acids.^39–42^

Unexpectedly, we observed a decline in 3OC12-HSL levels in the supernatant in *lasI*-expressing cells after 24 hours (Fig. 1d). We examined if generalized toxicity or metabolic burden could partially explain the decreasing 3OC12-HSL levels, however, the growth rate of *luxI*-expressing cells mimicked that of WT (Supplemental Fig. S2), indicating that AHL levels are in decline when growth rates begin to slow yet population density is high. Conversely, 3OC6-HSL accumulation remained relatively constant over time, with no significant changes in concentration after the initial rise. The reduced abundance of AHLs after 24 hours suggests that there may be intrinsic limitations on AHL synthesis. Cells entering stationary phase may have reduced AHL accumulation as has been shown previously in *Pseudomonas*^43^, alternatively nutrients or metabolic pre-cursors may also become limiting in late phase cultures^44^.

### 2.1. AHLs exhibit intrinsic lability and partitioning to biomass which limit extracellular accumulation

It is established that lactolysis of the lactone ring can contribute to inherent instability of AHLs^45,46^, while some organisms encode lactonases or other AHL-degrading enzymes. We therefore evaluated the contribution of natural AHL breakdown under laboratory growth conditions as compared to any *S. elongatus*-specific AHL degrading activities. We therefore evaluated the stability of chemically-defined AHLs (4000 nM AHL) in sterile BG11 and in paired flasks inoculated with *S. elongatus* (Fig. 2). Measurable levels of AHLs in the supernatant declined slightly faster in flasks containing *S. elongatus* (∼24 hours half-life) compared to sterile media (∼30 hours half-life; Figs. 2a, 2b). Both the length of the acyl chain (6C vs 12C) and illumination of the culture flasks did not significantly impact AHL stability (Figs. 2a, 2b). Separately, we were unable to identify any known homologs for lactonases, acylases, or other AHL-degrading enzymes that are encoded in the genome of *S. elongatus.*In the absence of a dedicated enzymatic path for AHL breakdown, we hypothesized that AHLs may partially partition to cell biomass and therefore be unavailable for extraction in the aqueous supernatant medium. We therefore setup an induction experiment, inducing either *lasI* or *luxI* with variable levels of IPTG (0.05 and 0.5 mM for *lasI* and 0.5 and 1.0 mM for *luxI*) as before, but concomitantly extracted AHLs from both the supernatant (Fig. 2c) and cell biomass (Fig. 2d). LC-MS analysis revealed that a significant amount of 3OC12-HSL was indeed present within the cell biomass (Fig. 2d). The raw quantity of AHL associated with cell biomass was similar to that found in the supernatant (Figs. 2c, 2d). However, the effective volume for cells is significantly lower than that of the supernatant, therefore the effective AHL concentration in biomass is substantially higher (∼37 to 80-fold: Table 1).Our results are generally consistent with prior reports that 3OC12-HSL preferentially accumulates in *P. aeroginosa* cell biomass, where it was estimated that ∼25% of the AHL partitions to the cell membranes^47^, possibly due to the chemical resemblance to fatty acids^48,49^. In some organisms, efflux pumps are used to promote secretion of AHLs from the cells^47^ although they do not appear to be necessary to enable sufficient AHL export in *S. elongatus*.

**Figure 2.**
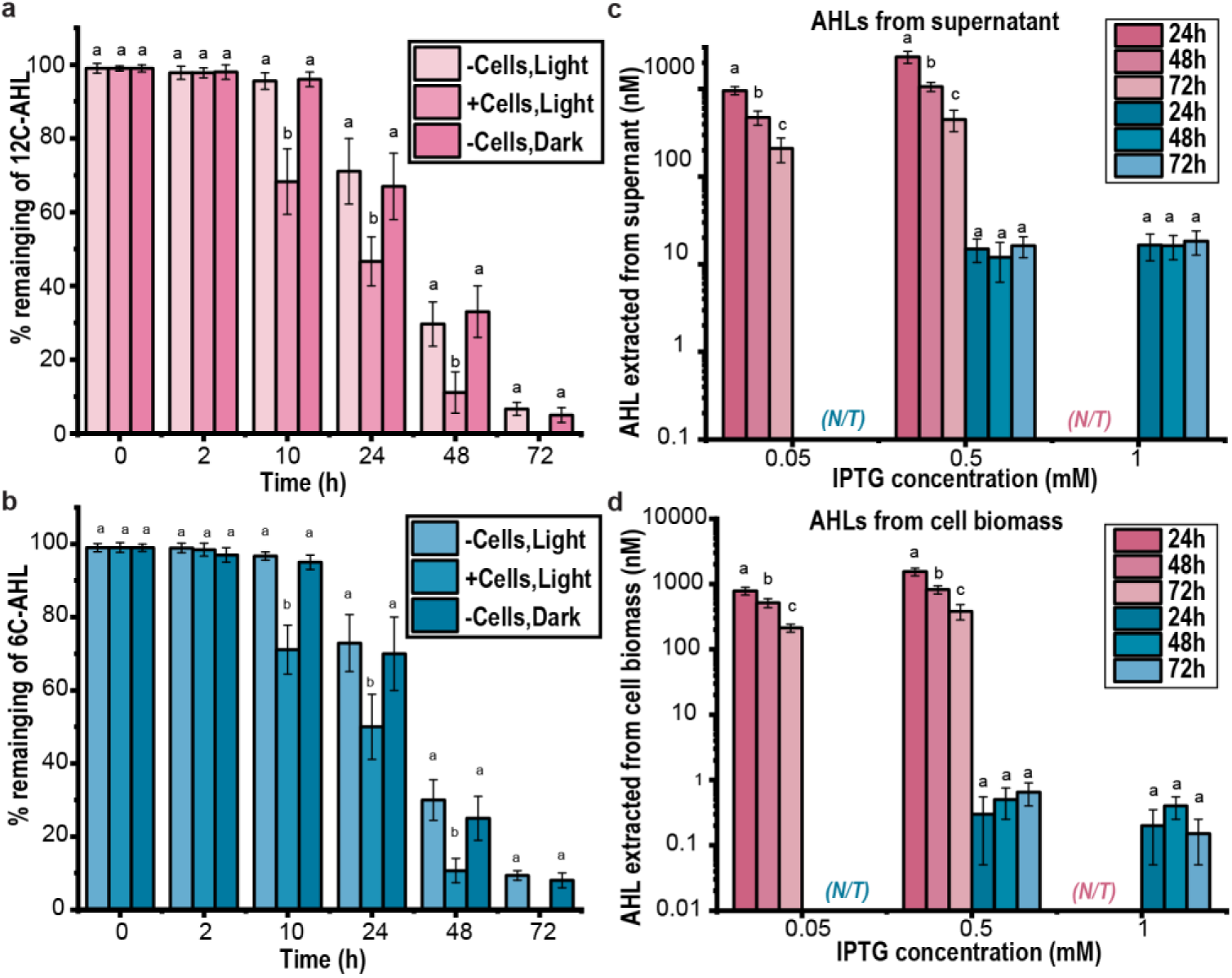
AHLs are labile and sequestered in *S. elongatus* cell biomass. Monitoring the abundance of a defined quantity of added 3OC12-HSL (a) or 3OC6-HSL (b) over time in sterile media or with *S. elongatus* cells under conditions of cellular growth (*i.e.*, illumination) or in the dark. Comparison of AHLs extracted from supernatant (c) and from *S. elongatus* biomass (d) over time in *S. elongatus* strains expressing LasI (pink) or LuxI with varied IPTG inducer concentrations. (N/T) indicates a “not tested” inducer concentration. Averages of ≥3 independent biological replicates are shown ± SD. Significance was calculated by one-way ANOVA followed by Tukey’s multiple comparison test. Bars labeled with different letters are significantly different (*P* < 0.05).

**Table 1.** Calculations of AHL accumulation in biomass relative to secreted compounds.

| Number of cells | Volume of spent media (mL) | Total volume of cells (converted to μL) | Ratio of volumes | Time point (hours) | Strain and IPTG concentration used (mM) |
| --- | --- | --- | --- | --- | --- |
| $1.554 \times 10^9$ | 0.5 | 6.33 | 79 | 24 | LasI 0.5 |
| $2.562 \times 10^9$ | 0.5 | 10.36 | 48 | 48 | LasI 0.5 |
| $3.330 \times 10^9$ | 0.5 | 13.52 | 37 | 72 | LasI 0.5 |
| $1.518 \times 10^9$ | 0.5 | 6.16 | 81 | 24 | LuxI 1.0 |
| $2.643 \times 10^9$ | 0.5 | 10.72 | 47 | 48 | LuxI 1.0 |
| $3.237 \times 10^9$ | 0.5 | 13.14 | 38 | 72 | LuxI 1.0 |

Taking together, our data suggests minimal enzymatic modification of AHL molecules by *S. elongatus*. Given documentation that AHL breakdown can be accelerated under conditions of elevated temperature (above 22°C) or pH (>7.2)^46,50^, and prior reports that AHLs breakdown substantially in LB over 30 hours^51^, we suggest that abiotic factors are the dominant contributor to the AHL loss over time we observe over time (Fig. 2). Free AHL levels may also appear artificially deflated due to preferential proportioning of these small, hydrophobic molecules to cell biomass. These parameters collectively determine the available pool of functional AHLs to drive engineered quorum sensing pathways.

It is established that lactolysis of the lactone ring can contribute to inherent instability of AHLs^45,46^, while some organisms encode lactonases or other AHL-degrading enzymes. We therefore evaluated the contribution of natural AHL breakdown under laboratory growth conditions as compared to any *S. elongatus*-specific AHL degrading activities. We therefore evaluated the stability of chemically-defined AHLs (4000 nM AHL) in sterile BG11 and in paired flasks inoculated with *S. elongatus* (Fig. 2). Measurable levels of AHLs in the supernatant declined slightly faster in flasks containing *S. elongatus* (∼24 hours half-life) compared to sterile media (∼30 hours half-life; Figs. 2a, 2b). Both the length of the acyl chain (6C vs 12C) and illumination of the culture flasks did not significantly impact AHL stability (Figs. 2a, 2b). Separately, we were unable to identify any known homologs for lactonases, acylases, or other AHL-degrading enzymes that are encoded in the genome of *S. elongatus.*In the absence of a dedicated enzymatic path for AHL breakdown, we hypothesized that AHLs may partially partition to cell biomass and therefore be unavailable for extraction in the aqueous supernatant medium. We therefore setup an induction experiment, inducing either *lasI* or *luxI* with variable levels of IPTG (0.05 and 0.5 mM for *lasI* and 0.5 and 1.0 mM for *luxI*) as before, but concomitantly extracted AHLs from both the supernatant (Fig. 2c) and cell biomass (Fig. 2d). LC-MS analysis revealed that a significant amount of 3OC12-HSL was indeed present within the cell biomass (Fig. 2d). The raw quantity of AHL associated with cell biomass was similar to that found in the supernatant (Figs. 2c, 2d). However, the effective volume for cells is significantly lower than that of the supernatant, therefore the effective AHL concentration in biomass is substantially higher (∼37 to 80-fold: Table 1).Our results are generally consistent with prior reports that 3OC12-HSL preferentially accumulates in *P. aeroginosa* cell biomass, where it was estimated that ∼25% of the AHL partitions to the cell membranes^47^, possibly due to the chemical resemblance to fatty acids^48,49^. In some organisms, efflux pumps are used to promote secretion of AHLs from the cells^47^ although they do not appear to be necessary to enable sufficient AHL export in *S. elongatus*.

Taking together, our data suggests minimal enzymatic modification of AHL molecules by *S. elongatus*. Given documentation that AHL breakdown can be accelerated under conditions of elevated temperature (above 22°C) or pH (>7.2)^46,50^, and prior reports that AHLs breakdown substantially in LB over 30 hours^51^, we suggest that abiotic factors are the dominant contributor to the AHL loss over time we observe over time (Fig. 2). Free AHL levels may also appear artificially deflated due to preferential proportioning of these small, hydrophobic molecules to cell biomass. These parameters collectively determine the available pool of functional AHLs to drive engineered quorum sensing pathways.

### 2.3 Combining AHL secretion and sensing circuits allows for time- and population-dependent gene activation in *S. elongatus*

We next sought to characterize the output for combined AHL secretion and sensing circuits in *S. elongatus*. We recently have shown that *S. elongatus* expressing either *luxR* or *lasR* are capable of responding to exogenously supplied AHLs by activating gene expression under respective promoters ^36^. As AHLs could be produced via the expression of *luxI* or *lasI*, we therefore constructed full QS circuits in which AHL molecules were both produced and sensed by the same population of cells. To achieve this, we co-expressed the AHL synthetase (*e.g.*, *lasI* or *luxI*) along with its cognate transcription factor (*e.g.*, *lasR* or *luxR*), both under IPTG-inducible promoters (Figs. 3a and 3d). The inducible promoter allowed convenient characterization of how tuning the expression levels of these genes impacts the circuit output. We encoded a fluorescent reporter under the control of the cognate promoter (P*_lux_*_I_/P*_lasI_*) to provide a functional readout of QS circuit activation, and monitored cellular fluorescence over time.

**Figure 3:**
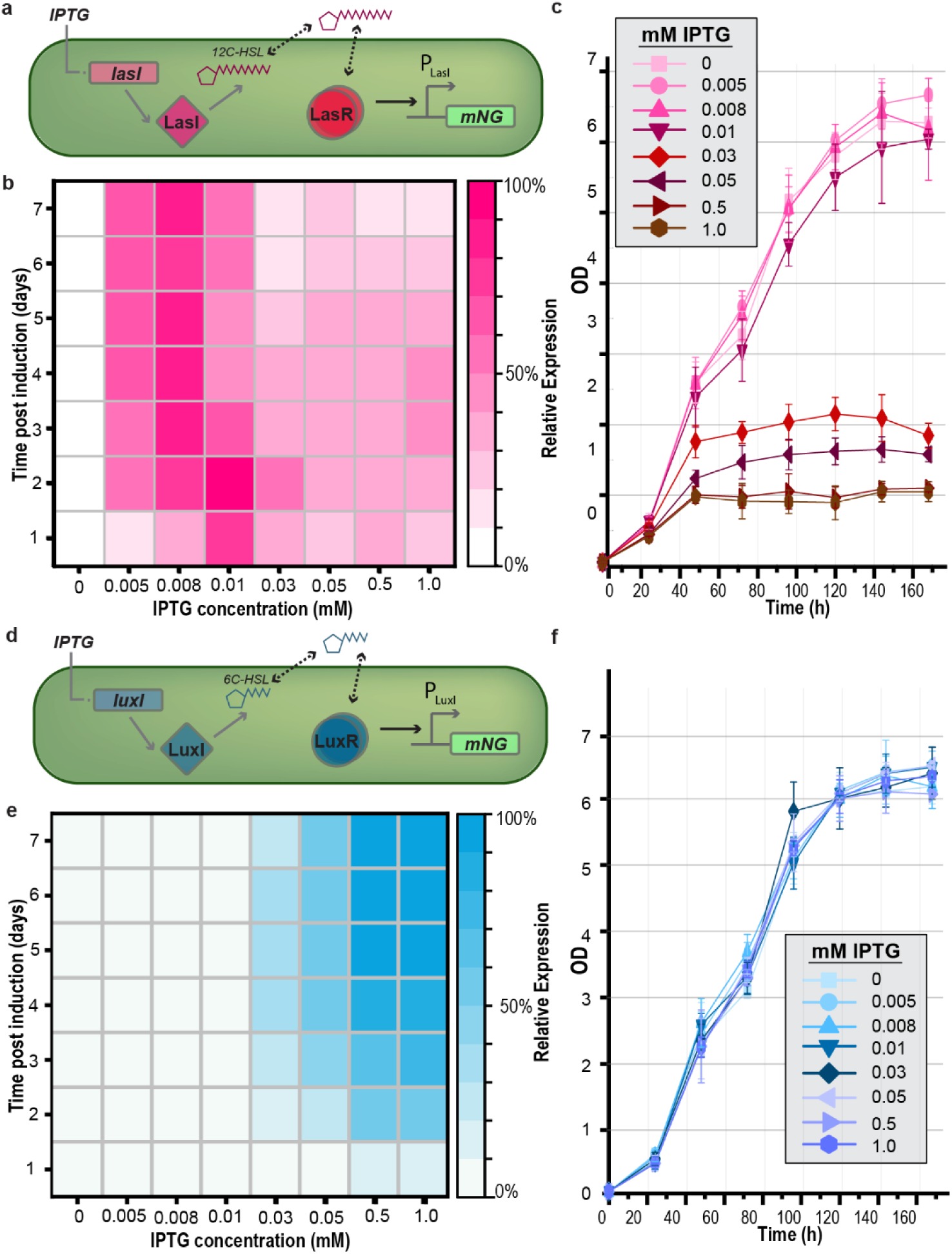
Characterization of LasR/LasI and LuxR/LuxI QS circuits. Overview diagram illustrating the quorum sensing-based genetic circuit design of LasR/LasI (a) and LuxR/LuxI (d). IPTG controls the expression of the AHL synthetase gene *lasI(*a,pink)*/luxI*(d,blue) and the transcription regulators LasR/LuxR. Output of the QS circuits are monitored over time as a function of expression of the fluorescent reporter mNG, which is transcribed from a promoter bound by LasR (b) or LuxI (d), respectively. Reporter output is normalized to the highest intensity over a 7 day experiment with varied [IPTG] Growth curves measuring OD750 of *S. elongatus* expressing the LasR/lasI (c) or LuxR/luxI (e) QS circuits under various IPTG concentrations. Averages of ≥3 independent biological replicates are shown ± SD.

#### 2.3.1 Effectiveness of the LasR/lasI circuit is limited by toxicity of LasR

We observed unexpected fluorescence output from *S. elongatus* strains co-expressing LasI and LasR, with reporter expression suppressed at higher levels of induction and at later points of the culture growth phase (Figs. 3a, 3b, 3c). Specifically, higher fluorescence output was highest under minimal IPTG induction (≤0.01 mM), where our LC/MS data indicated only a minimal amount of 3OC12-HSL is produced (20 nM; Fig. 1d). Above 0.01 mM IPTG induction, fluorescence reporter levels were decreased and showed additional declines as the culture grew (Fig. 1d, Supplemental Figs. S4b, S4d). Cultures with higher induction also displayed signs of general toxicity, as measured by OD (Fig. 3c) and visible chlorosis (Supplemental Fig. S4c), suggesting a more general toxicity of the circuit. It has been previously shown that LasR can cause a significant growth impediment in *P. aeruginosa* and *E. coli*,^36,52–55^ therefore, our results suggest LasR may have inherently cytotoxic properties when expressed at higher levels and/or stabilized by the cognate binding of 3OC12-HSL. The significant growth arrest in *S. elongatus* and failure to show late-phase activation in dense cultures indicates that LasR/lasI does not fit the criteria for a suitable design of a cyanobacterial QS circuit, although it does display high sensitivity at low-to-intermediate induction conditions.

#### 2.3.2 Despite low AHL secretion rates, LuxR/luxI circuits allow for QS-like activation patterns

By contrast, reporter output for strains with *luxI* and *luxR* genes was consistent with increasing pathway activation with increased cell density (Fig. 3d, 3e, Supplemental Fig. S4). *S. elongatus* cultures underwent ≥60-fold increase in cell density over the week-long growth curve, saturating around day 5 (Fig. 3f). We observed no significant impact on growth or cell pigmentation at higher induction levels (Fig. 3f, Supplemental Fig. S5c). Instead, there was a strong correlation between induction and reporter output, and pathway output generally increased at higher cell densities (Fig. 3e, Supplemental Figure S5b). However, the *luxR/luxI* circuit was much less sensitive than *lasR/lasI* (Fig. 3b, e), an observation that is likely tied to the relatively low production levels of 3OC6-HSL, which are roughly two orders of magnitude lower than 3OC12-HSL production under most conditions measured (Fig. 1e). FACS analysis revealed that the increase in reporter output from activated luxR/luxI circuits was consistent across all cells of the population, rather than an increase in a subpopulation of cells (Supplemental Fig. S5e). In total, the *luxR/luxI* circuit displayed a QS-like pattern of late-phase activation, reaching output levels approximately 5-8-fold above paired uninduced controls.

#### 2.3.3 Crosstalk between Lux and Las enables construction of an improved, hybrid cyanobacterial circuit

LuxR-family receivers display known cross-reactivity to “off-target” AHLs, likely a function of non-specific binding of AHLs with longer or shorter acyl chains to the hydrophobic pocket in the N-terminus ^53,56^. Generally, LuxR-family receivers display a preference for a specific chain length, but also bind other AHLs with less sensitivity (within 1-3 orders of magnitude^54,57^). We therefore examined if the high LasI AHL production rates could be combined with the non-toxic LuxR-sensing module to produce a hybrid pathway with improved QS functions (Fig. 4a). *S. elongatus* cells expressing a *luxR/lasI* hybrid circuit indeed exhibited output that was directly related to both induction level and stage of the cell culture (Figs. 4a, 4b). Compared to the Lux-only circuit, the hybrid circuit exhibited improved sensitivity and was responsive to induction levels as low as 0.01 mM IPTG (Fig. 4b). Additionally, no growth defects or chlorosis of the cells expressing the luxR/lasI circuit were evident, even at high induction levels and dense cultures (Fig. 4c and Supplemental Fig. S6d).

**Figure 4.**
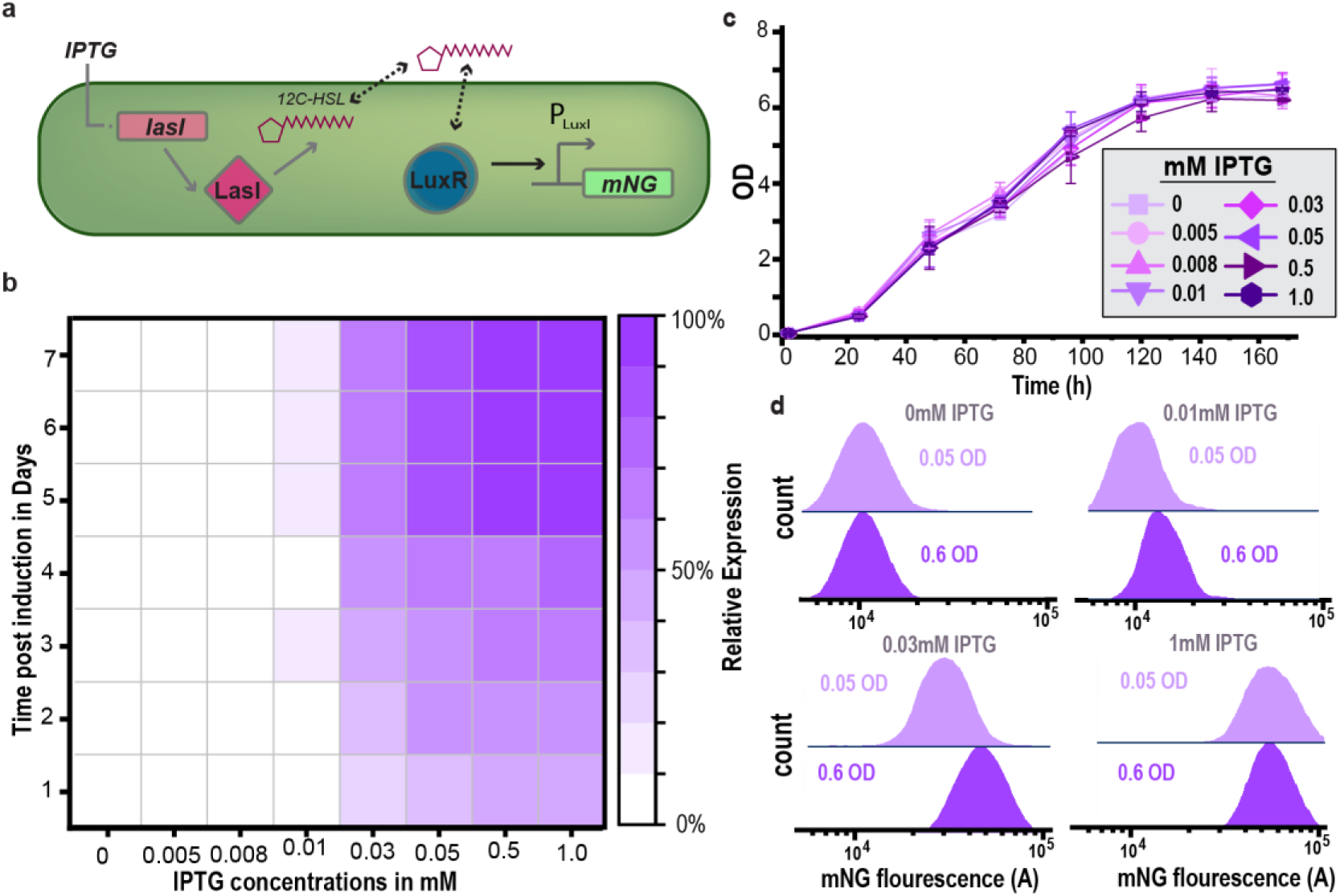
Circuit characterization of LuxR/LasI QS. Overview diagram illustrating the hybrid QS circuit design of LuxR/LasI. IPTG controls the expression of the AHL synthetase gene lasI(pink) and the transcription regulators LuxR (blue). In the presence of sufficient AHL LuxR is stablized and binds the PLuxI promoter to drive mNG reporter expression (a). Heatmap of QS mNG signal output normalized to the highest recorded intensity over 7 day experiments with varied [IPTG] (b). Growth curves measuring OD750 of *S. elongatus* expressing the hybrid LuxR/lasI QS circuit under various IPTG concentrations. Averages of ≥3 independent biological replicates are shown + SD (c) cultures. Representative density histograms of mNG reporter output from cultures initiated with low (0.05 OD) or high (0.6 OD) culture densities at 24 hours post induction (d).

While the hybrid *lasR*/*luxI* circuit had the highest output in late phase cultures, we could not be certain if output expression was tied to population density, as in canonical QS systems, or was merely a function of increasing output over time. We therefore varied the starting concentration of cells and followed the fluorescence reporter activation over time using a range of inducers. At low concentrations of inducer driving the expression of the AHL synthetase (0.01 and 0.03 mM of IPTG), we observed that the starting biomass did impact reporter output. However, at high inducer concentrations (1mM IPTG) output expression was not significantly different between the low density and high-density cultures, suggesting that cells synthesizing a high level of AHLs (Fig 1d) may “self-activate” by retention/sensing of AHLs in the cytosol before they equilibrate with the surrounding medium (Fig. 4d).

### 2.4 Comparison of LasR/LasI, LuxR/LuxI and hybrid LuxR/LasI circuit performance

We compared the performance of the three QS circuits described above: LasR/LasI, LuxR/LuxI, and a hybrid system (LuxR/LasI) over seven days. To better compare the utility of these pathways for potential applications, we represent the activation of the circuit as a fold-change in output expression at varied IPTG concentrations as compared to the uninduced reference (Fig. 5, Supplemental Fig. S7). The LasR/LasI system showed high responsiveness to low IPTG concentrations from day one, but its performance declined at higher concentrations over time due to toxic impacts on cell growth and chlorosis. The LuxR/LuxI system exhibited delayed activation, only responding to the higher IPTG concentrations (≥ 0.03 mM) and with a limited tunability (between 0.03 and 0.5 mM). The hybrid system demonstrated improved sensitivity (≥ 0.01 mM) and tunability (0.01-1mM). The hybrid *luxR/lasI* circuit also reached a higher fold induction under the optimal configurations (10-fold activation), maintaining reporter output levels under activating conditions up until 7 days (Fig. 5).

**Figure 5.**
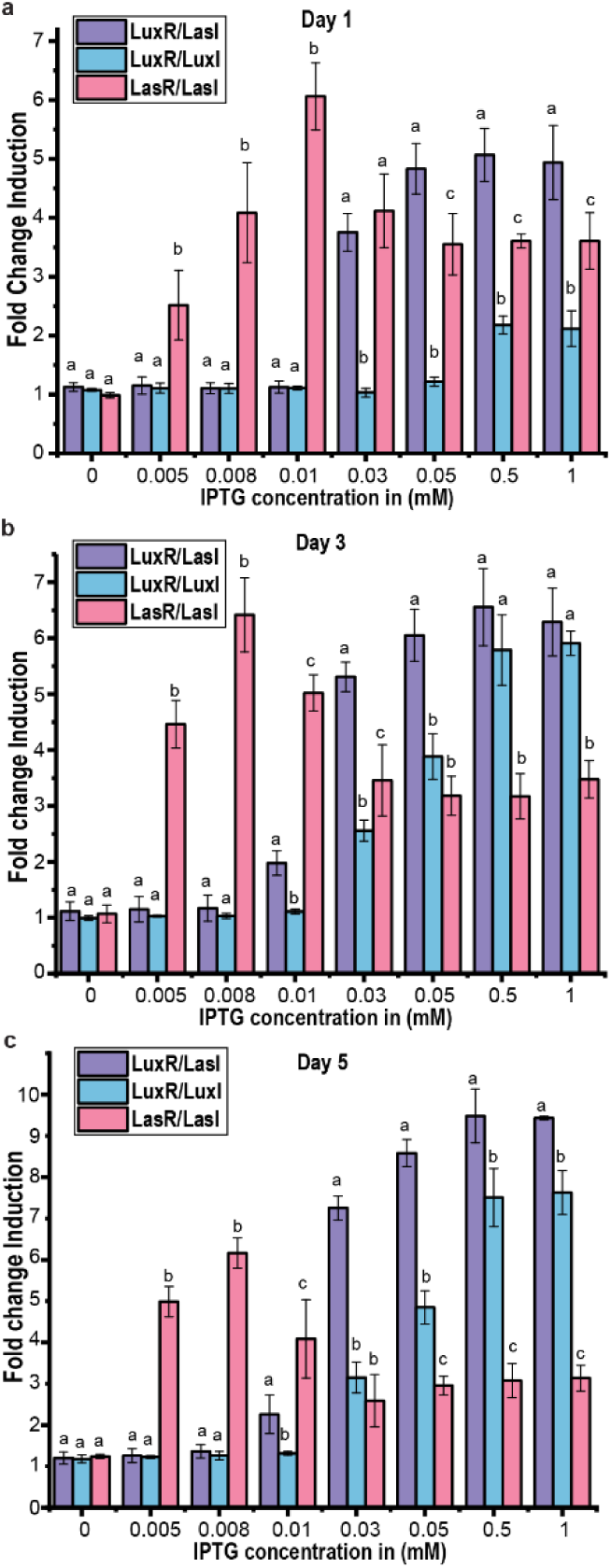
Comparison of different QS circuit outputs. The fold change of the reporter output for the three QS circuits (LuxR/LuxI; blue, LasR/LasI; pink, and LuxR/LasI; purple) as calculated by the maximum value divided by the basal (uninduced) mNG expression at the indicated IPTG concentrations. Data are reported for cultures 1 Day (a), 3 days (b) and 5 days after induction. Averages of ≥3 independent biological replicates are shown + SD. Significance was calculated by one-way ANOVA followed by Tukey’s multiple comparison test. Bars labeled with different letters are significantly different (P < 0.05).

Unexpectedly, despite the decline in measurable AHL levels in the supernatant of late-phase cultures, we did not observe a decline in reporter output for either the *luxR/luxI* or *luxR/lasI* circuits (Fig. 5, *e.g.*, Day 7). In native QS pathways, there is often a positive feedback element encoded that acts to increase the ‘switch-like’ behavior for pathway activation and can contribute to hysteresis whereby the pathway exhibits resistance to transient fluctuations of AHL – effectively "remembering" the prior activation state^53,58^.However, in these engineered circuits, there is no inbuilt feedback regulation to maintain an activated circuit output. Possible explanations for the maintenance of activation could include: i) reduced protein dilution rates in late-phase cultures due to decreased growth rates; or; ii) a residual pool of AHL molecules that are maintained in the biomass phase even when a decreased level of AHLs are available in the supernatant medium (Fig. 2; Table 1).

### 2.5 Proof of concept QS circuit improving biomass harvesting in late-phase cultures

The introduced *luxR/luxI* and *luxR/lasI* QS circuits exhibited favorable properties for activation of genetic pathways that could be programmed for self-activation in late-phase cyanobacterial cultures. One current problem associated with large scale cultivation of cyanobacteria is that the energy inputs required to harvest and dewater cell biomass can be a significant contributor to the total production costs^31,32^. While the morphological features of filamentous cells can improve cell recovery, many cyanobacterial models with the best genetic toolkits do not have filamentous growth modes. Furthermore, filamenting cells may require additional energy input during the growth phase to maintain adequate mixing in cell suspensions. We therefore sought to link a gene controlling cell morphology to the output of a QS circuit to create a pathway that induced cells to adopt a filamentous morphology only at a late growth phase.

Toward this goal, we placed the *cdv3* gene fused with GFP under the control of the P*_luxI_* promoter (Supplemental Fig. S8A). Cdv3, is a protein localized to midzone of dividing *S. elongatus* cells that has homology to the DivIVA division regulator in *Bacillus subtilis* and is involved in FtsZ-ring closure. Misregulation of Cdv3 has been shown to result in cell hyper elongation, as cells continue to grow but fail to form a divisome that can contract to bisect the cell ^59–62^. As expected, P*_luxI_*::*cdv3* bearing cells which also contain a copy of the LuxR transcriptional regulator were able to respond to the addition of exogenous 3OC6-HSL by inducing cell division arrest and increasing cell lengths, as viewed by fluorescence microscopy (Supplemental Fig. S8).

We next evaluated linking the Cdv3-based cell elongation phenotype to the previously described *luxR/lasI* or *luxR/luxI* heterologous QS circuits (Fig. 6a). Engagement of the hybrid *luxR/lasI* system drove dramatic cell elongation starting as early as 24 hours and continuing to increase average cell length through day 3 (Fig. 6b, 6c Supplemental Figs. S9, S10). Cells maintained a hyperelongated phenotype (>50-100 µm) through 5 days, the longest time point that we measured (Supplemental Fig. S10). By contrast, *cdv3* expression driven by the *luxR/luxI* based circuit required a higher IPTG induction, yet resulted in a milder elongation phenotype, as viewed by microscopy and FACS (Fig. 6b, Supplemental Figs. S11, S12). The reduced capacity of *luxR/luxI* to drive hyperelongation is consistent with the lower maximal induction of this circuit relative to the hybrid QS design (Supplemental Figs. S5, S6). Relatedly, under very strong IPTG induction, *luxR/lasI* hybrid cells at higher density also exhibited variability in the chl *a* distribution within hyperelongated cells (Supplemental Fig. S9), a phenotype consistent with prior evidence in cells with strongly upregulated Cdv3 levels^59^. To verify that the elongation phenotype was due to intercellular signaling (*i.e.*,not self-activation) we incubated two separate strains together; 1) the sender strain bearing only the AHL synthetase and; 2) the receiver LuxR-Cdv3/GFP. Confirming that the AHL signal is able to equilibrate from the producing cells to the medium and activate the receivers, we see two cellular morphologies in these mixed experiments: WT-length and elongated subpopulations (Supplemental Figs. S13 and S14).

**Figure 6:**
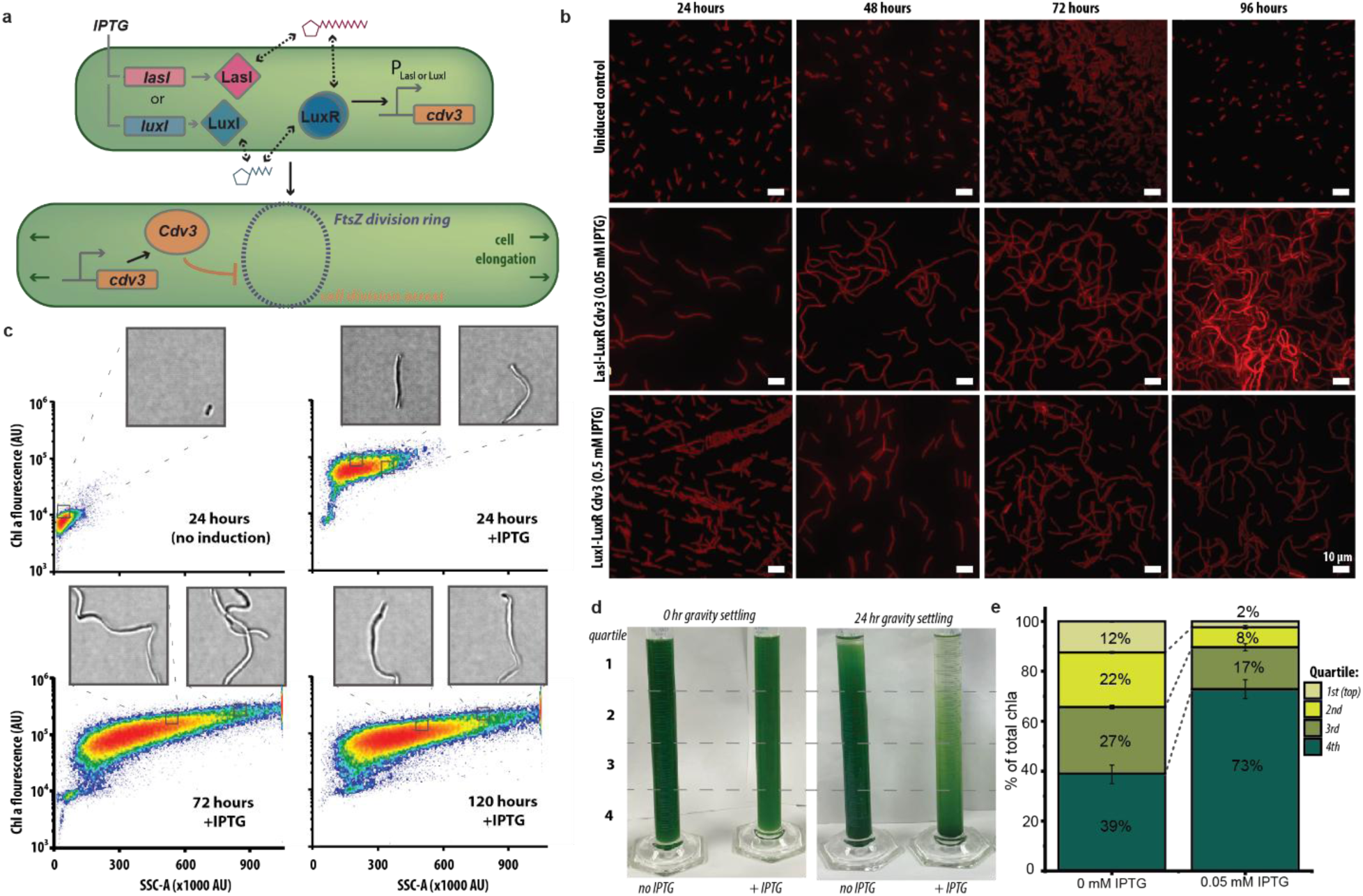
QS circuits drive a auto-inducing cell elongation pathway to improve biomass recovery in late-phase S. elongatus cultures. Cartoon schematic of LuxR/Cdv3-GFP/LasI and LuxR/Cdv3-GFP/LuxI strains. Detection of secreted AHL molecules drives expression of the cell division factor, *cdv3*, inducing cell elongation by preventing cell fission. b) Representative images of elongating cells from 24-96 hours following induction of the hybrid LuxR/Cdv3-GFP/LasI or LuxR/Cdv3-GFP/LuxI QS circuits. Cells are visualized by autofluorescence of chlorophyll *a* (red). Scale bar: 10 μm. c) Elongation of the hybrid LuxR/Cdv3-GFP/LasI induced at 0.05 mM of IPTG strain impacts the entire population of cells, as monitored by FACS analysis over time. Inset brightfield images are representative cells within the population that were imaged in real-time on the Attune flow cytometer. Chla-A; area of chlorophyll a; SSC-A, side scatter area. d) Hyperelongated cells are more rapidly sedimented under normal gravity, as demonstrated by images of LuxR/Cdv3-GFP/LasI cultures transferred to graduated cylinders after 3 days of induction (0.05 mM IPTG) ; left image 0-hour gravitation settling, right image after 24 hours of gravitation settling. e) Quantification of the fraction of cells in each quadrant of the water column after 24 hours gravity sedimentation, as measured by Chl*a* content. Averages of ≥3 independent biological replicates are shown ± SD

To assess the unassisted rate of sedimentation under gravity, we induced *luxR/lasI* cultures with 0.05 mM IPTG and grew them for 72 hours before transferring liquid cultures to graduated cylinders and imaging sedimentation over time. The hyperelongated cells from induced hybrid cultures visually sedimented faster relative to uninduced controls (Fig. 6d). We quantified the gravity-based sedimentation rate after 24 hours by fractionation of the culture and extracted chl *a* to evaluate the proportion of cells contained in each quadrant of the water column (Fig. 6e). The cell elongation caused by self-signaling from the hybrid QS circuit was improved relative to the addition of exogenous AHLs to the same circuit (Supplemental Fig. S8c, S8d).

## 3. Discussion

Herein, we have evaluated three variants of a QS circuit for their capacity to both produce sufficient AHL molecules and to accumulate to levels sufficient to autoinduce linked genes in cyanobacteria. We show that cyanobacteria can synthesize and respond to AHL at physiologically-relevant levels such that components of the LuxR/luxI family can be utilized to construct QS systems for late gene activation in dense cyanobacterial cultures. We characterized the kinetics of cyanobacterial QS-linked genetic outputs as a function of a range of different expression levels for the AHL-synthesis and AHL-receiving proteins. In a proof of concept circuit, we show that features of cyanobacterial cell morphology can be programmed to become more amenable for harvesting in late-phase, higher-density cultures. Taken together, our design showcases potential applications of QS-based genetic circuits for bioproduction in cyanobacteria.

While our proof of concept focuses upon late phase activation of a genetic module that improves biomass recovery by changing cell morphology, a variety of possible circuits could be implemented for the purposes of enabling cyanobacterial bioproduction. Most straightforwardly, QS systems could be used for late-phase activation of metabolic pathways to enhance target bioproduct synthesis at later phases of culture density, potentially mitigating fitness losses during phases of rapid cell growth in culture ^63,64^. Other cellular features could also be tied to late-phase activation, such as pathways to improve bioproduct recovery through decreasing cell wall integrity or inducing cell lysis: lytic genes derived from cyanophages that degrade the cell wall might be utilized for product release in such a strategy ^65^. Using repressive systems or NOT logic gates, it would be feasible to invert the QS output to turn down native pathways at later phases of cell growth or to keep a genetic pathway active only during the early phases of culture growth. For example, light penetration into dense cyanobacterial cultures is often suboptimal due to expansion of light-harvesting antenna that led to absorption of the majority of light in only the uppermost layer of the light column. A variety of antenna truncating strategies have been reported in the literature ^66–68^ which could be used to suppress antenna size or refine antenna properties in late phase cultures via QS-based approaches. Flexible genetic pathways for suppression of target genes, such as CRISPRi or protein degradation tags^69,70^, could be used to deactivate native or engineered pathways upon reaching a critical population density in this manner.

In this context, we find that features of the three tested QS circuits (LuxR/luxI, LasR/lasI, and LuxR/lasI) are distinct in ways that may influence their utility for driving downstream genes of interest. Overall, we observe that the hybrid system – utilizing LasI for AHL production and LuxR for AHL detection – is the most robust, combining features of the high production of 3OC12-HSL without the negative cytotoxic components of LasR activity. The hybrid system drives the highest stable expression of QS-linked genetic outputs over time and exhibits the greatest ON:OFF ratio from the maximal induction relative to the basal expression level. Perhaps due to the lower productivity of 3OC6-HSL, the LuxR/luxI QS circuit activates with slower kinetics and at later phases of the growth curve (Fig. 5). By either using less active AHL synthetases (*e.g.*, LuxI) or expressing them under weaker promoters, genetic circuits desirable to activate only near the end of the logarithmic growth phase could be designed in cyanobacteria. By contrast, the LasR/lasI circuit is compromised by the apparent toxicity of accumulating too high of a cellular level of the LasR transcriptional regulator. However, the LasR/LasI system responds to very low concentrations of 3OC12-HSL and activates at relatively lower expression and intermediate cell densities. Therefore, the LasR/LasI system could be suitable for controlling pathways that would be ideal to express during the middle stages of logarithmic growth, although any realistic applications would likely need to mitigate the cell toxicity of the LasR protein, perhaps through the construction of a hybrid 3OC12-HSL–responsive receptor with improved compatibility.

Curiously, we note that the level of AHLs available in the supernatant becomes limited at later growth phases, likely due to a complex interplay of inherent instability of AHL molecules and reduced per-cell AHL secretion rates in later culture phases. Encoding a positive feedback loop similar in nature to those observed in many native QS pathways might be appropriate to further tune the output features of engineered circuits, particularly if more ‘switch-like’ activation kinetics are desirable. Somewhat surprisingly, we find that outputs for both a fluorescent reporter and the Cdv3-elongation are maintained at a high level of activation in late phase cultures, despite the declining supernatant AHL abundance. It is possible that this may be attributed to preferential retention of AHLs in the biomass fraction or to relatively slow turnover of proteins in slower-growing cells at later culture phases. In this context, we see that driving expression of the AHL synthetase family member at too high of a level can lead to self-activation that becomes more unlinked from the culture density (Fig. 4d). A lower basal expression of AHL synthetase likely allows for equilibration of the autoinducing signal between the producing cell and the surrounding medium. Alternatively, exploration of AHL export pathways, such as the *mexA-mexB-oprM*-encoded efflux pump from *Pseudomonas aeruginosa*, could help to increase the rate of AHL secretion from producing cells^47^. Additional research on these features would assist in the design of more complex QS-based cyanobacterial circuits.

Properly designed, QS systems could be used to bypass the need for expensive inducible compounds that are not economically-feasible in large scale cultivation. We note that the QS based circuits that we feature in this work are still tied to tunable P*_trc_* promoters that require IPTG to tune the expression level of the LuxI and LuxR family components. While this was a convenient approach for systematically exploring the impact of expression level on the features of QS pathway output, any scaled application would require the identification of suitable constitutive promoter elements that can drive gene expression within a desired range. The target strength of such constitutive promoters would be dependent upon the demands for the particular application in question. Finally, we note that signals that can readily diffuse between cells, like the family of AHL molecules, could be used to distribute genetic circuits across more than one partner species in engineered microbial communities – an area currently of increased scientific interest in the field^36,71–74^.

## 4. Methods

### 4.1 Strains and culture conditions

*S. elongatus* cultures were grown in BG11 medium supplemented with 1 g L^−1^ HEPES to a final pH of 7.7 with NaOH. Flasks were cultured in a Multitron incubator (Infors HT) at 32°C under ambient air CO_2_ or supplemented with 2% CO_2_ with ∼150 μmol photons m^−2^s^−1^ of light provided by Sylvania 15 W Gro-Lux fluorescent bulbs and shaken at 150 rpm. Cultures were back-diluted daily to an OD_750_ of 0.3 and acclimated to the medium/irradiance for at least 3 days prior to experiments. Chloramphenicol (Cm; 25 μg mL^−1^) and Kanamycin (Kn; 12.5 μg mL^−1^) was used to maintain LuxR/mNG/LuxI, LasR/mNG/LasI, LuxR/mNG/LasI,LuxR LuxR/Cdv3-GFP/LasI and LuxR/Cdv3-GFP/LuxI containing cells. Kanamycin (Kn; 12.5 μg mL^−1^) was used to maintain the LuxI and LasI. Chloramphenicol (Cm; 25 μg mL^−1^) was used to maintain the LuxR/Cdv3-GFP. In all cases, antibiotic selection was removed prior to conducting the reported experiments to minimize unintended effects. All strains used in this study are listed in Table 2.

**Table 2.** Cyanobacterial strains used in this study.

| Strain | Relevant genotype and phenotype<sup>a</sup> | Plasmids used to generate this strain | Source or reference<sup>b</sup> |
| --- | --- | --- | --- |
| *S. elongatus* PCC 7942 |  |  | PCC |
| LasI | *S. elongatus* PCC 7942 with *P<sub>trc</sub>::luxI* integrated at NS2; Kn<sup>r</sup> | p1047 | This work |
| LuxI | *S. elongatus* PCC 7942 with *P<sub>trc</sub>::luxI* integrated at NS2; Kn<sup>r</sup> | p1048 | This work |
| LuxR/mNG/LuxI | *S. elongatus* PCC 7942 with *P<sub>trc</sub>::luxR_P<sub>luxI</sub>::mNG* integrated at NS3, and with *P<sub>trc</sub>::luxI* integrated at NS2; Cm<sup>r</sup> Kn<sup>r</sup> | p1048 and p1013 | 36 |
| LasR/mNG/LasI | *S. elongatus* PCC 7942 with *P<sub>trc</sub>::lasR_P<sub>lasI</sub>::mNG*; integrated at NS3, and with *P<sub>trc</sub>::luxI* integrated at NS2; Cm<sup>r</sup> Kn<sup>r</sup> | p1047 and p1016 | 36 |
| LuxR/mNG/LasI | *S. elongatus* PCC 7942 with *P<sub>trc</sub>::luxR_P<sub>luxI</sub>::mNG*; integrated at NS3, and with *P<sub>trc</sub>::luxI* integrated at NS2; Cm<sup>r</sup> Kn<sup>r</sup> | p1048 and p1016 | 36 |
| LuxR/Cdv3-GFP | *S. elongatus* PCC 7942 with *P<sub>trc</sub>::luxR_P<sub>luxI</sub>::cdv3* integrated at NS3; Cm<sup>r</sup> | p1051 | This work |
| LuxR/Cdv3-GFP/LasI | *S. elongatus* PCC 7942 with *P<sub>trc</sub>::luxR_P<sub>luxI</sub>::cdv3* integrated at NS3 and with *P<sub>trc</sub>::lasI* integrated at NS2; Cm<sup>r</sup> Kn<sup>r</sup> | p1051 and p1047 | This work |
| LuxR/Cdv3-GFP/LuxI | *S. elongatus* PCC 7942 with *P<sub>trc</sub>::luxR_P<sub>luxI</sub>::cdv3* integrated at NS3 and with *P<sub>trc</sub>::luxI* integrated at NS2; Cm<sup>r</sup> Kn<sup>r</sup> | p1051 and p1048 | This work |

<sup>a</sup>Kn<sup>r</sup>, kanamycin resistance; Cm<sup>r</sup>, chloramphenicol resistance; NS2, neutral site 2; NS3, neutral site 3.
<sup>b</sup>PCC, Pasteur Culture Collection.

### 4.2 Strain construction

Plasmids 1013, 1016 and 1051 were generated using isothermal assembly from either PCR-amplified or synthesized dsDNA (Integrated DNA Technologies, IDT), whereas 1047 and 1048 plasmids were generated using restriction enzyme cloning^75^. Constructions for integration of DNA in the cyanobacterial genome were flanked with 300–500 bp of homology to promote efficient homologous recombination^76^. Coding sequences were codon-optimized for *Synechocystis* sp. PCC 6803 using the IDT codon optimization tool. *S. elongatus* cells were transformed as previously described^77^. Chemically competent *E. coli* DH5α were prepared and transformed as routine^78^. All constructs were confirmed by PCR and Sanger sequencing. Plasmid details are reported in Table 3. Plasmids 1047,1048 and 1051 are submitted to Addgene and are awaiting assignment of an accession number, we will finalize this information in any revision of this manuscript.

**Table 3.** Plasmids used in this study.

| Plasmid | Relevant genotype and phenotype<sup>a</sup> | Source or reference |
| --- | --- | --- |
| p1013 | pNS3_ *P<sub>trc</sub>::luxR_P<sub>luxI</sub>::mNG*; Cm<sup>r</sup> | 36 |
| p1016 | pNS3_ *P<sub>trc</sub>::lasR_P<sub>lasI</sub>::mNG*; Cm<sup>r</sup> | 36 |
| p1051 | pNS3_ *P<sub>trc</sub>::luxR_P<sub>luxI</sub>::cdv3:GFP*; Cm<sup>r</sup> | This work |
| p1047 | pNS2_ *P<sub>trc</sub>::lasI*; Kn<sup>r</sup> | This work |
| p1048 | pNS2_ *P<sub>trc</sub>::luxI*; Kn<sup>r</sup> | This work |

<sup>a</sup>Cm<sup>r</sup>, chloramphenicol resistance; Kn<sup>r</sup>, kanamycin resistance.

### 4.3 AHL Extraction and LC–MS Analysis

Cyanobacterial cultures were typically seeded at OD_750_ = 0.3 with the indicated IPTG concentrations (from 0.001 mM up to 2 mM). Cells were pelleted by centrifugation of 0.5 mL samples at 13,300 *xg* for 10 min, and the supernatant was kept for the extraction. To quantify the AHLs in the supernatant, 10 μL of 5 μM N-hexanoyl-homoserinelactone-d_3_ (N_6_-HSL-d_3_) was used as an internal standard and was added with 1 mL of ethyl acetate acidified as the extraction solvent. 0.5 mL of the extraction solvent and 0.5 mL supernatant was vortex-mixed for 1 min, centrifuged for 1 min and the organic phase (top) was collected. The extraction procedure was repeated, and the ethyl acetate volume extracts (1 mL total) were dried using a Speedvac vacuum concentrator (Thermo Scientific). The samples were reconstituted in 100 μL of 50% methanol/water immediately prior to LC– MS/MS analysis, resulting a final concentration of the internal standard of 0.5 μM. For whole cell extraction, 3 mL of cyanobacterial culture was collected in a cellulose filer paper via pressure vacuum. The cells were washed with 3 mL of H_2_O to ensure that any residual AHLs were removed. and the filer papers were cut into small pieces and were placed in 1.5 mL eppendorf tube with 1 mL of 80% MetOH with 0.5 μΜ internal standard. An overnight extraction was performed in -20°C followed by 20 min of 250 rpm shaking under dark. The liquid phase was collected and passed through a MetOH filter to remove residual cell biomass.

Metabolite extracts were analyzed on a Thermo Q-Exactive UHPLC LC–MS/MS system (Thermo Electron North America). The mobile phase was 0.1% formic acid in Milli-Q water (A) and acetonitrile containing 0.1% formic acid (B). The stationary phase was an Acquity reverse phase UPLC BEH C-18 column (2.1 mm × 100 mm, Waters, Milford, MA). The chromatographic run was as follows: 0–0.5 min hold at 2% B, ramp to 40% B at 1 min, ramp to 99% B from 1 to 7 min, hold at 99% B until 8 min, return to 2% B at 8.1 min, and hold at 2% B until 10 min. The injection volume was 10 μL, the flow rate was 0.30 mL/min– 1, and the column temperature was at 40 °C. Mass spectra were collected using positive mode electrospray ionization with a full MS/AIF (all ion fragmentation) method with a scan range set from m/z 80 to 1200. The capillary voltage was 3.5 kV, transfer capillary temperature was 256.25 C, sheath gas was set to 47.5, auxiliary gas was set to 11.25, the probe heater was at 412.5 C, and the S-lens RF level was set to 50. The MS mode and AIF scans were acquired at 70,000 resolution with an AGC target of 3 × 106 (maximum inject time of 200 ms), and the stepped normalized collision energies for the AIF scans were 10, 20, and 40. Raw files (.raw) were analyzed using the Thermo Xcalibur software. For quantification purposes, standard curves of each AHL molecule quantified were created in the range of 0.96–6000 nM using the same internal standard as mentioned above. The ratios of LC–MS/MS peak areas of the analyte/internal standard were calculated and used to construct calibration curves of the peak area ratio against the analyte concentration using unweighted linear regression analysis. For the calibration curve standard AHLs were commercially available: 3-oxohexanoyl-l-homoserine lactone (3OC6-HSL, K3007, Sigma-Aldrich) for the lux system, N-(3-oxododecanoyl)-l-homoserine lactone (3OC12-HSL, O9139, Sigma-Aldrich) for the las system.

To estimate internal AHL concentration, we measured AHL from whole cell extractions as above and estimated the cell volume. This calculation involved utilizing a standard curve that correlates OD_750_ to cell number (Supplemental Fig. 3) that the cell number was calculated using an attune flow cytometer.

Average cell width and length measurements were calculated for the cyanobacterial strains based on fluorescence microscopy. Total cell volume was calculated by multiplying the total number of cells with the volume calculated from a cell of average dimensions using the bellow equation^79^.

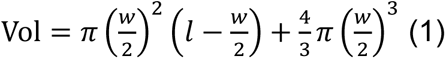

### 4.4 Flow Cytometry

All flow cytometry measurements were performed in live cells on a 4-laser Attune CytPix with a CytKick Max Autosampler Software 6.2.0. *S. elongatus* cultures were diluted in BG11 media to reach OD_750_ = 0.5 OD baseline, followed by a 1:10 dilution in FACS buffer (1× PBS, 2% (v/v) BSA, 2 mM EDTA, 2 mM sodium azide). The following optical configuration was used for each fluorophore (excitation|emission): mNeonGreen [488nm|530/30], GFP [488nm|530/30], Chlorophyll [405nm|660/20] (486V). These settings were maintained for all species-specific experiments of each AHL family (*i.e.*, Lux and Las), ensuring that the relative expression levels between systems could be compared. Cyanobacterial samples were gated using FSC (400V) and SSC (400V) to distinguish the singlet population and with the chlorophyll to remove debris and noise, more than 5,000 cells were measured per sample. The raw data were analyzed using the FCS Express 7 software (De Novo Software, USA). For each sample, the median fluorescence intensity (MFI) was calculated. Mean and standard deviation were calculated from independent biological replicates. The induction ratio for each circuit was calculated as the maximal MFI divided by the basal MFI.

### 4.5 Microscopy

All live-cell microscopy was performed on cells in exponential growth by centrifuging 2 mL of culture at 10,000 × g for 5 min, resuspending into 80 μL of BG11, and transferring a 2 μL aliquot to a 3% agarose pad. The cells were allowed to briefly equilibrate and be absorbed by the agarose (≥10 min) before the pad was placed onto a #1.5 glass coverslip for imaging. Images were captured using a Zeiss Axio Observer D1 inverted microscope equipped with an Axiocam 503 mono camera and a Zeiss Plan Apochromat 63X 1.4 NA oil-immersion lens.

### 4.6 Sedimentation assays

Samples were back-diluted into BG-11 medium every 24 hours for up to 72 hours in 50-ml flask volumes. Cultures were induced with the indicated IPTG or 3OC6-HSL and grown for 3 days under standard growth conditions. 25 mL of culture was transferred to 25-mL graduated cylinders and allowed to settle by gravity for 24 hours on a benchtop. To quantify settling, 4 equal volume fractions (6.25 mL each) were pipetted carefully from the top phase. 1 mL of cell culture from each section was used for the chlorophyll *a* (chl *a*) extraction protocol and to accurately measure the sedimentation process.

### 4.7 Chlorophyll *a* extraction

We adapted the established protocol from^80^ to measure chlorophyll *content*. Briefly chl *a* was extracted from 1 mL of cyanobacteria pellet centrifuged at 16000 *xg* for 10 min and then resuspended in 1 mL 100% MeOH. Cells were extracted for 30 min at 4°C and vortexed at high-speed vortex for 2 min. Cells were pelleted and absorbance was taken at 665 nM to estimate chl *a* as previously published. ^81^

### 4.8 Statistical analysis

Recorded measurements are represented as mean values, with error bars expressing the SD of n≥3 biological replicates experiments, as indicated. The significance of differences between groups was evaluated by one-way ANOVA followed by Tukey’s multiple comparison test or by an unpaired Student’s t-test. Statistical analyses were carried out using GraphPad Prism software (GraphPad Software Inc., San Diego, CA). Differences were considered statistically significant at P < 0.05.

## 5. Conclusions

In this study we demonstrate that complete QS circuits can be successfully integrated into *S. elongatus* and couple the output of these circuits to induction of population-density dependent genetic outputs. We characterized three QS systems with different patterns of sensitivity and activation, demonstrating that two of these systems can drive late-phase activation of an engineered output in high-density cultures. By coupling QS systems with the *cdv3* gene, we achieved a dramatic cell elongation of *S. elongatus* cells in late-phase cultures, which leads to enhanced rates of cell sedimentation that could be utilized to improve cell biomass harvesting and dewatering. In total, our findings hold potential for industrial applications, particularly in addressing scalability challenges in cyanobacterial biotechnology. Future elaboration and enhancement of QS circuit design, such as incorporating positive or negative feedback loops and constitutive promoters, could further enable auto-induction systems with ideal properties for large-scale production.

## 6. Acknowledgments

The Attune Cytpix, located in the MSU Flow Cytometry Core Facility, is supported by the Equipment Grants Program, award no. 2022-70410-38419, from the U.S. Department of Agriculture (USDA) National Institute of Food and Agriculture (NIFA). We thank all the members of the Ducat lab for their help revising the article. We thank Dr. Tony Schilmiller for helping with LC–MS for AHL quantification.

## 7. Author contributions

E.J.K. and D.D. conceived the project. E.J.K., D.D., D.V. and M.S.M. designed the experiments, E.J.K. and S.G. performed the experiments, E.J.K and S.G. analyzed the data. E.J.K, D.C.D, M.S.M. wrote the manuscript and all authors reviewed, edited and approved the manuscript.

