## Supplemental S1-S14 for "Autoinduction expression systems via engineered quorum-sensing circuits in *Synechococcus elongatus PCC 7942*"

**Supplemental Material**

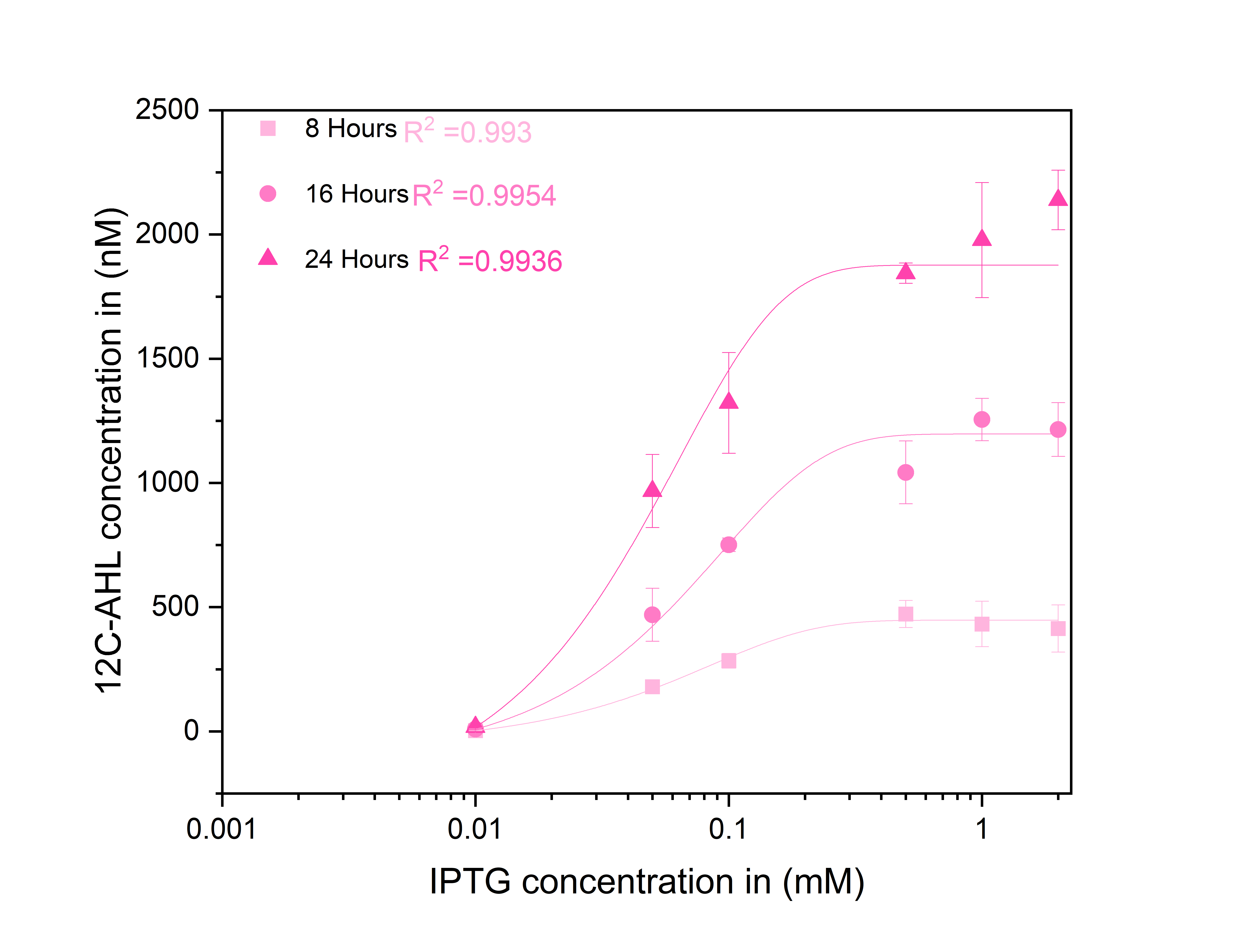

**Supplemental Figure S1.** **Dose response curves for 3OC12-HSL production**. The curves were measured over 24 hours at varied concentrations of IPTG induction of the *lasI* construct. Each line represents the relationship between the inducer and 3OC12-HSL secreted at one timepoint (8, 16 or 24 hours). Averages of ≥3 independent biological replicates are shown ± SD. The R-square (R^2^) values for the dose-response fitting of each candidate are displayed..

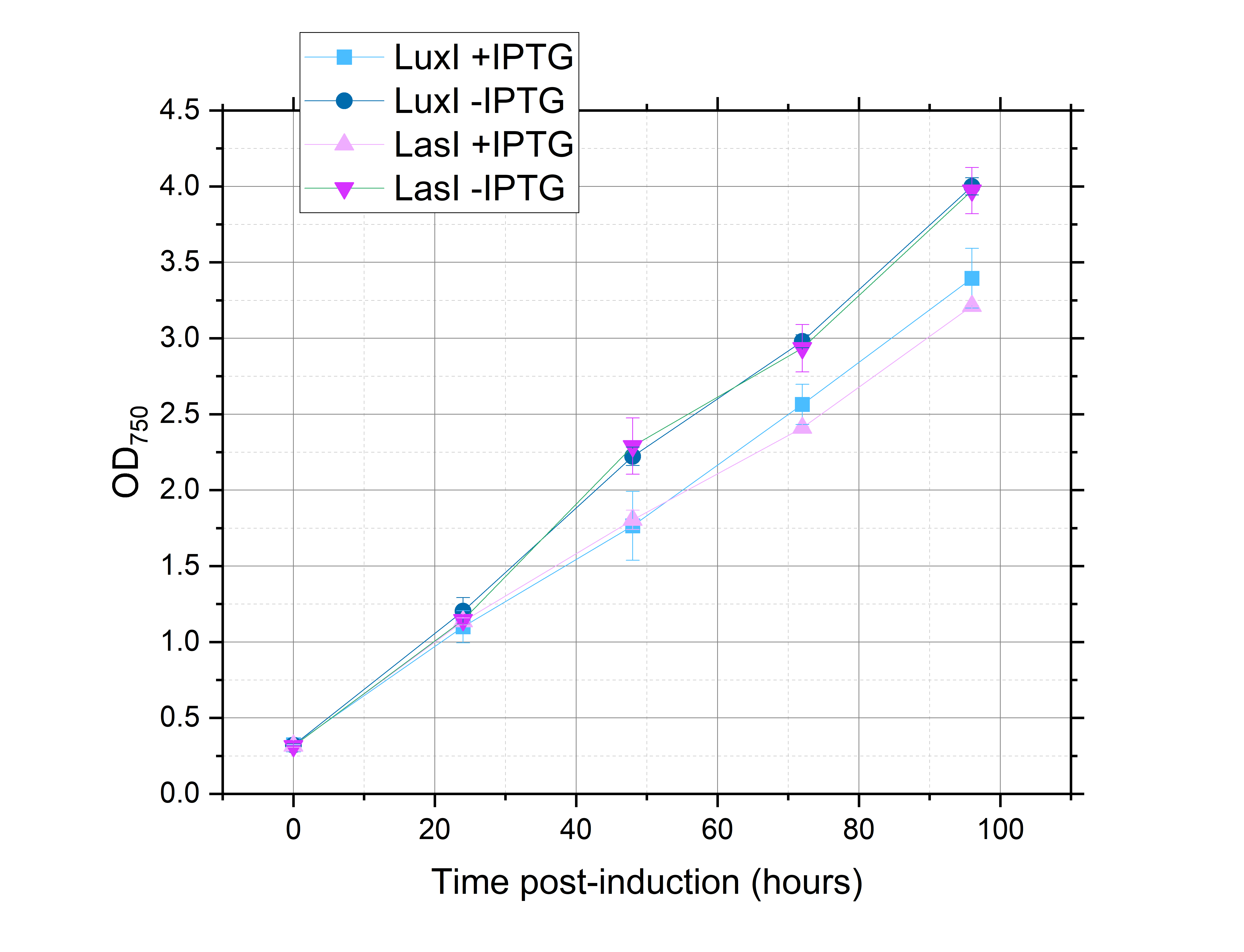

**Supplemental Figure S2.** **Growth curves of *S. elongatus* cultures**. The cultures were maximally induced for expression of LasI (0.5mM IPTG) and LuxI (1 mM IPTG) as compared to their corresponding uninduced controls. Averages of ≥3 independent biological replicates are shown ± SD.

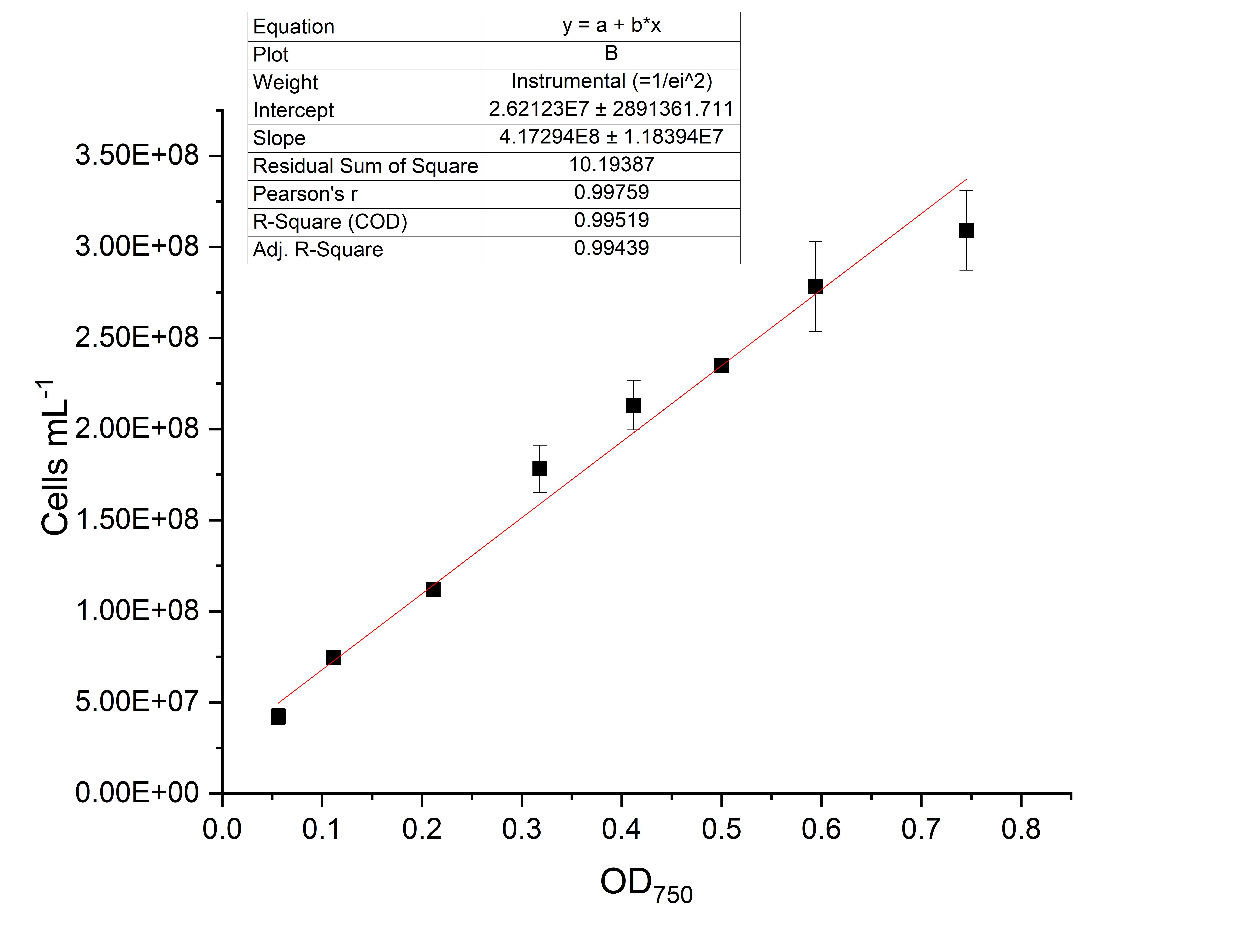

**Supplemental Figure S3. Standard curve correlating cells/mL with optical density.** The cells/mL were recorded using attune flow cytometer. Averages of ≥3 independent biological replicates are shown ± SD.

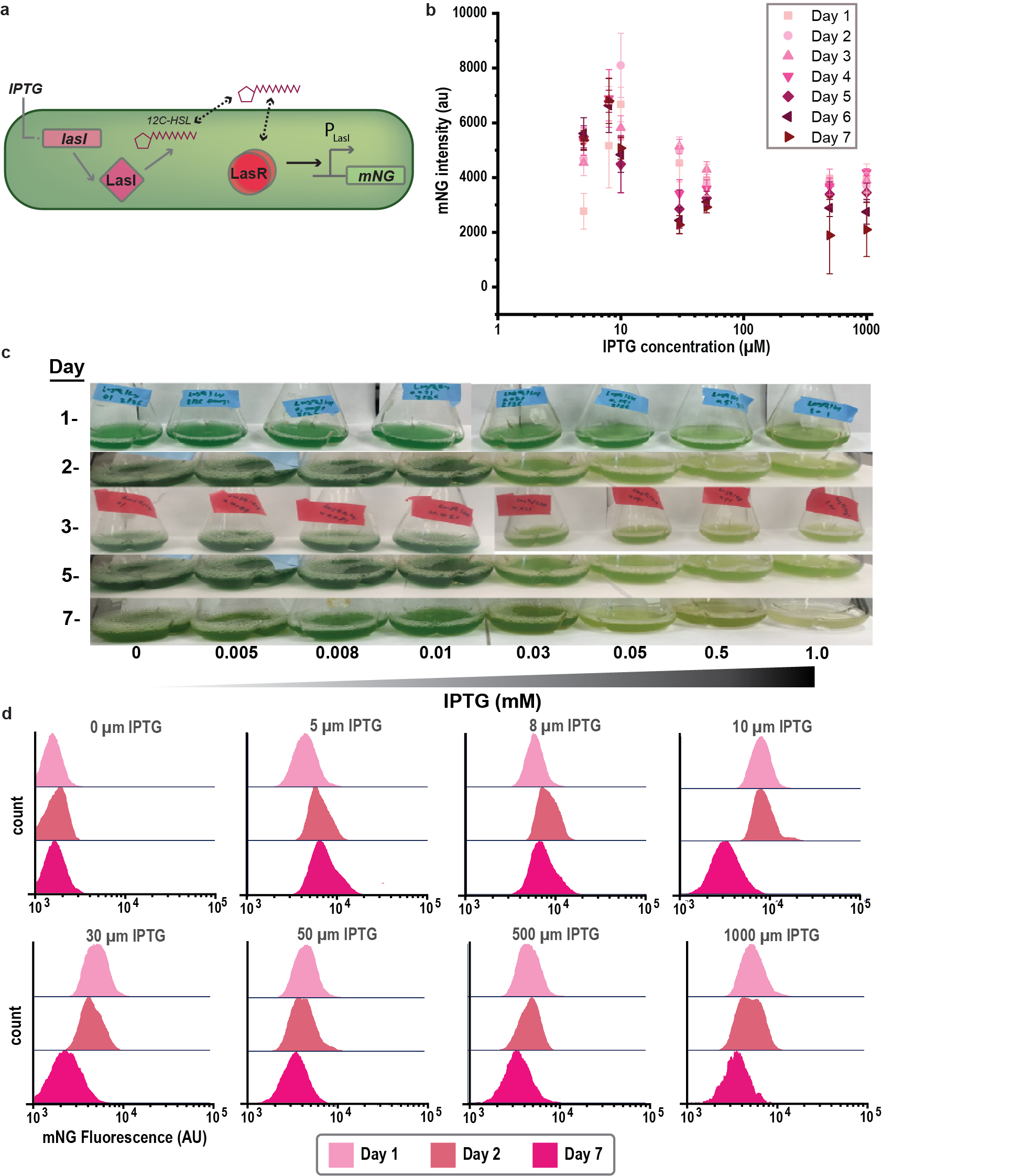

**Supplemental Figure S4.** **LasR/LasI system characterization.** Overview diagram illustrating the quorum sensing-based genetic circuit design LasR/LasI. IPTG controls the expression of the AHL synthetase gene *lasI* and the transcription regulators *lasR*, once enough AHL signal is sensed from the transcription regulator is dimerized and interacts with P*_luxI_* to turn the mNG fluorescent reporter. (a) Diagram showing the plotted values mNG values of the different IPTG concentration overtime and the error bars represent the standard deviation of 3 independent biological samples. Dose response curves were not able to be made due to the reduction of the signal. Averages of ≥3 independent biological replicates are shown ± SD.(b) Image showing the color of the culture flasks under an increase of IPTG concentration overtime. (c) Population histograms of the different IPTG concentrations for the days 1, 2 and 7 (the rest of the days were omitted for clarity) (d).

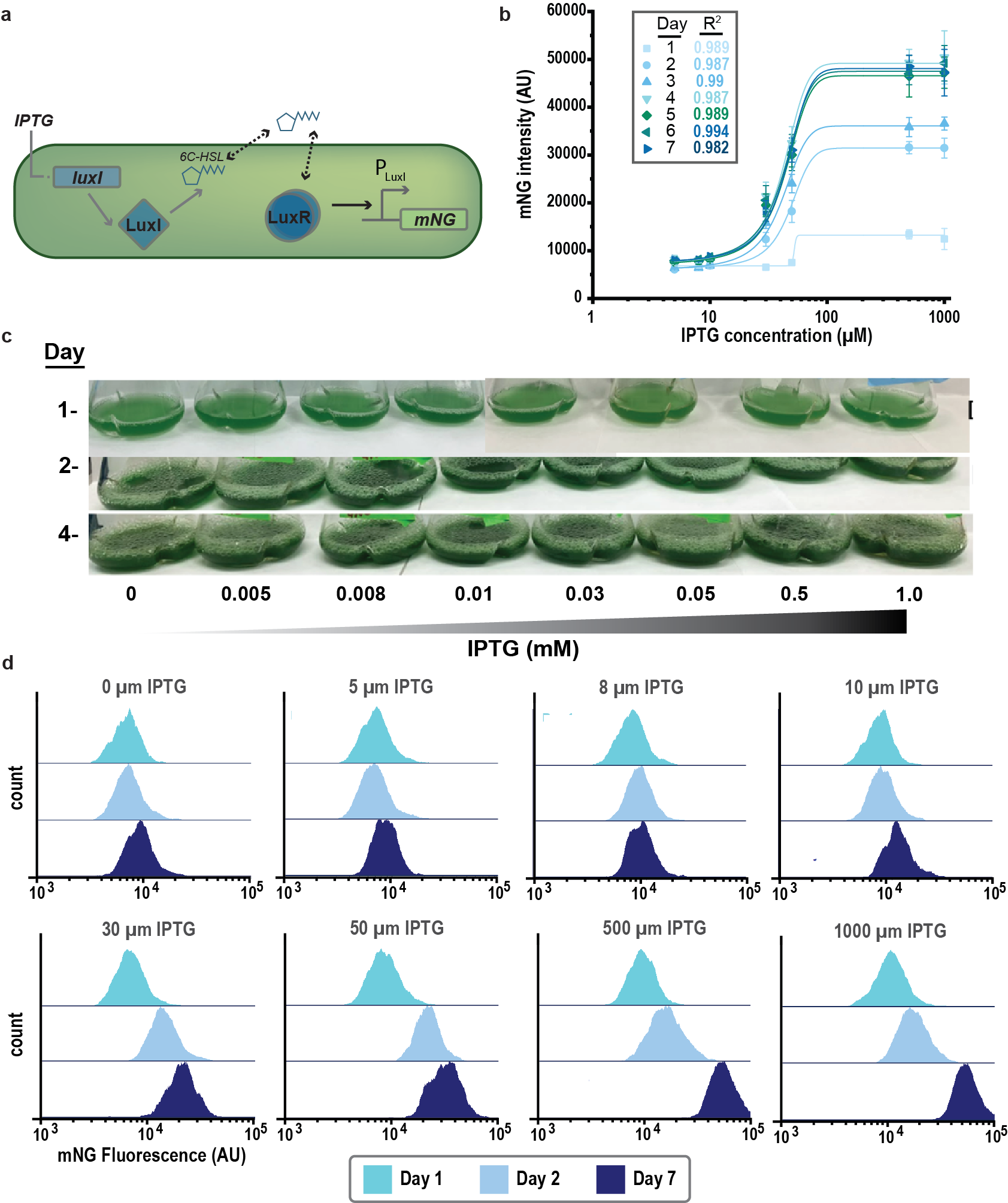

**Supplemental Figure S5.** **LuxR/LuxI system characterization.** (a) Overview diagram illustrating the quorum sensing-based genetic circuit design LuxR/LuxI. IPTG controls the expression of the AHL synthetase gene *luxI* and the transcription regulators *luxR*, once enough AHL signal is sensed from the transcription regulator is dimerized and interacts with P*_luxI_* to turn the mNG fluorescent reporter.(a) Diagram showing the plotted values mNG values of the different IPTG concentration overtime. Averages of ≥3 independent biological replicates are shown ± SD. The R-square (R^2^) values for the dose-response fitting of each candidate are displayed. (b) Images showing the color of the culture flasks under an increase of IPTG concentration overtime.(c) Population histograms of the different IPTG concentrations for the days 1, 2 and 7 (the rest of the days were omitted for clarity)(d) .

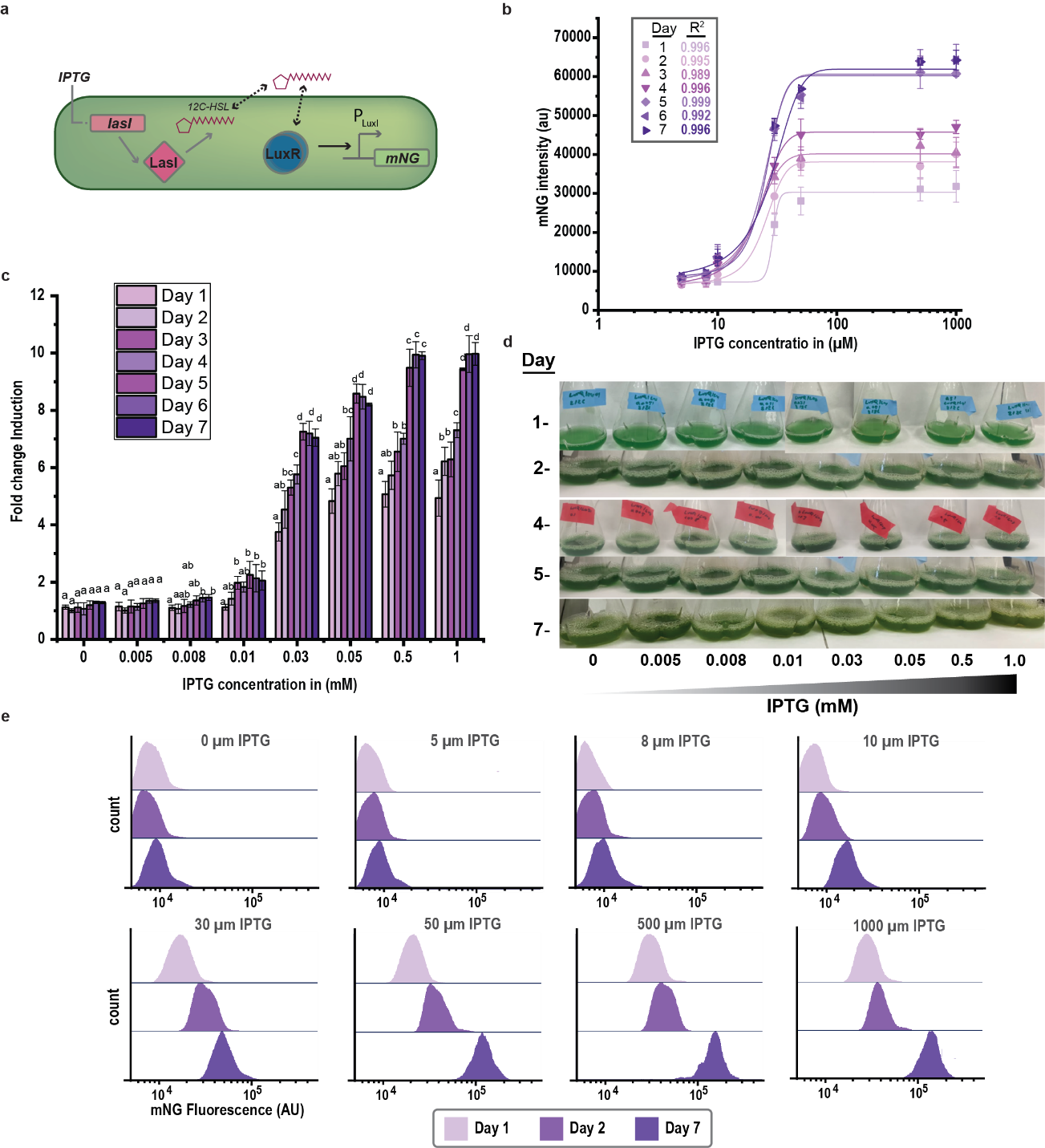

**Supplemental Figure S6**. **LuxR/LasI QS system characterization****.** Overview diagram illustrating the quorum sensing-based genetic circuit design LuxR/LasI . IPTG controls the expression of the AHL synthetase gene *lasI* and the transcription regulators *luxR*, once enough AHL signal is sensed from the transcription regulator is dimerized and interacts with P*_luxI_* to turn the mNG fluorescent reporter. (a) Diagram showing the plotted values mNG values of the different IPTG concentration overtime. Averages of ≥3 independent biological replicates are shown ± SD. The R-square (R^2^) values for the dose-response fitting of each candidate are displayed. (b) Fold-change induction of LuxR/LasI QS system overtime, IPTG used for induction from 0.005 to 1 mM of IPTG. The fold change induction of each QS circuit was calculated as calculated as the maximum over the minimum mNG intensity at specific IPTG concentrations relative to uninduced cognate controls. Averages of ≥3 independent biological replicates are shown ± SD. Significance was calculated by one-way ANOVA followed by Tukey’s multiple comparison test. Bars labeled with different letters are significantly different (P < 0.05). (c) Images showing the color of the culture flasks under an increase of IPTG concentration overtime. (d) Population histograms of the different IPTG concentrations for the days1, 2 and 7 (the rest of the days were omitted for clarity) (e).

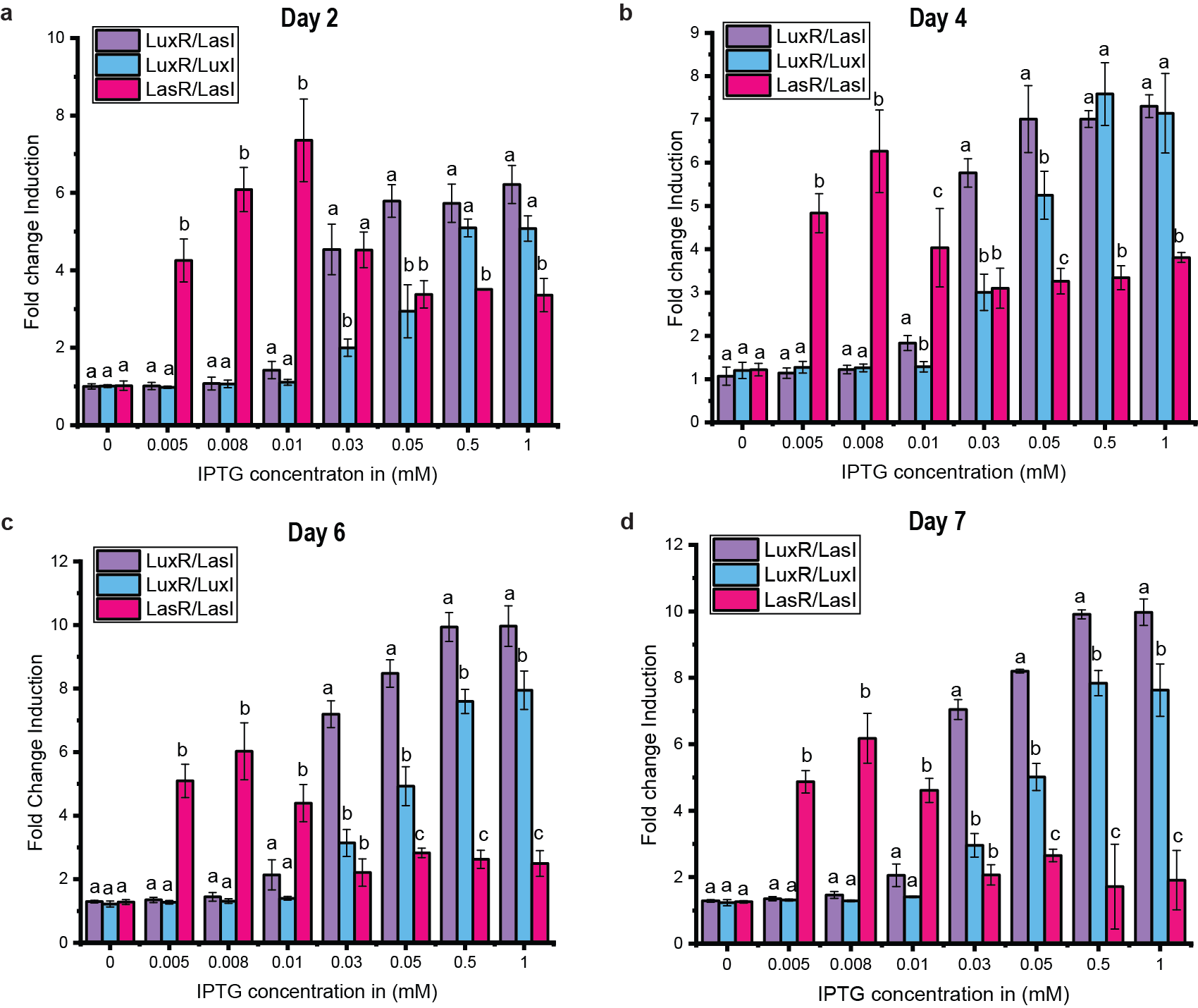

**Supplemental Figure S7. Fold change comparison of the QS cirquits LuxR/LuxI, LasR/LasI and LuxR/LasI overtime.** Fold change Induction comparison between the different IPTG concentrations overtime: (a) Day 2, (b) Day 4, (c) Day 6, and (d) Day 7. IPTG used for induction from 0.005 to 1 mM of IPTG. The fold change induction of each QS circuit was calculated as calculated as the maximum over the minimum mNG intensity at specific IPTG concentrations relative to uninduced cognate controls. Averages of ≥3 independent biological replicates are shown ± SD. Significance was calculated by one-way ANOVA followed by Tukey’s multiple comparison test. Bars labeled with different letters are significantly different (P < 0.05).

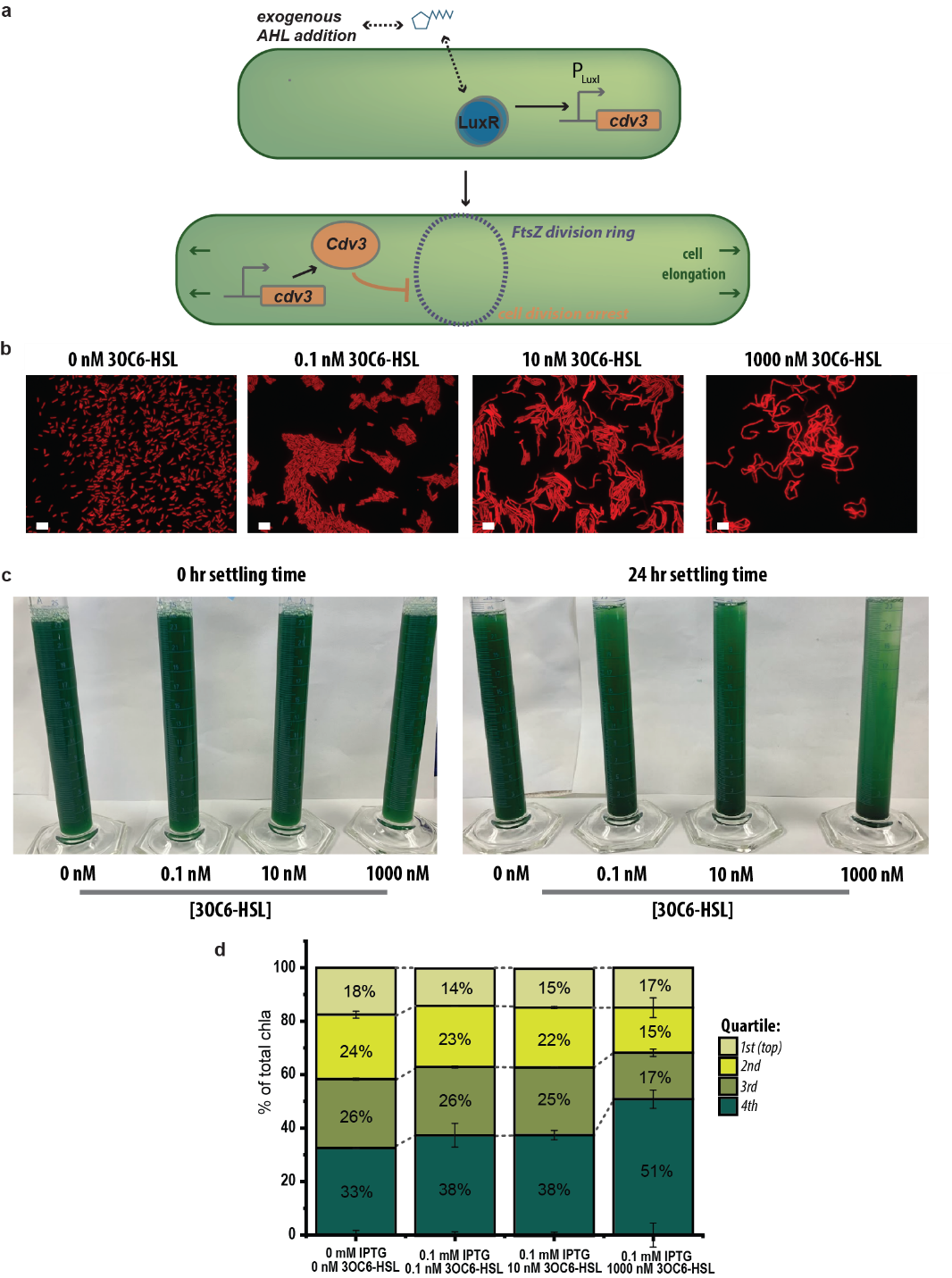

**Supplemental Figure S8. Characterization of the LuxR-Cdv3/GFP strain.** (a) Overview diagram illustrating the quorum sensing-based genetic circuit design LuxR-Cdv3/GFP. IPTG controls the expression of the transcription regulators *luxR*, once enough 3OC6-HSL (exogenously supplied) signal is sensed from the transcription regulator is dimerized and interact with Pl_uxI_ to turn the *cdv3* gene. The activated Cdv3 protein disrupts the formation of the FtsZ division ring, arresting cell division and making the cells elongate. Scale bar: 10μM.(b) Cell length monitor of the LuxR-Cdv3/GFP strain using a fluorescence microscopy under red channel (by tracking chl *a*) after three days of induction using 0.1 mM of IPTG and a titration range of exogenously supplied 3OC6-HSL signal. Scale bar: 10 μm. (c) Cell sedimentation assay after three days of 0.1 mM IPTG induction and a range of exogenously supplied 3OC6-HSL from 0.1 to 1000 nm. Left image shows the 4 different conditions after 0 hour of settling and right image shows the cyanobacteria settling after 24 hours. (d) Chl *a* quantification after 24 hours in different quarters of the cylinders across the various induction conditions normalized by the chl *a* at 0 hour settling. Averages of ≥3 independent biological replicates are shown ± SD.

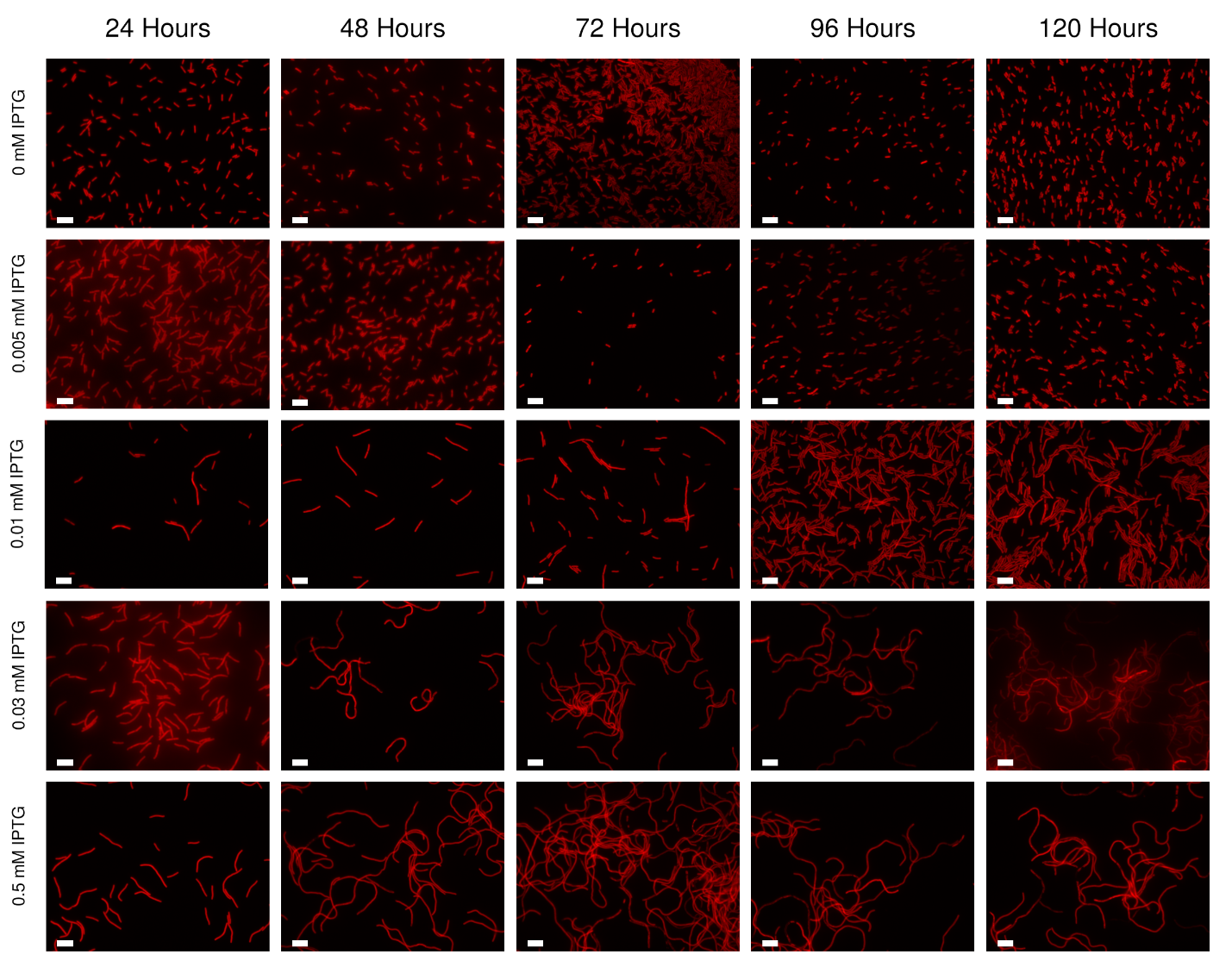

**Supplemental Figure S9. Characterization of the LuxR-Cdv3/GFP-LasI strain.** Cell length monitor overtime from 24 hours up to 120 hours of the LuxR-Cdv3/GFP-LasI strain and IPTG concentration ranging from 0 up to 0.5 mM, using a fluorescence microscope under red channel by tracking the chl *a*. Scale bar: 10 μm.

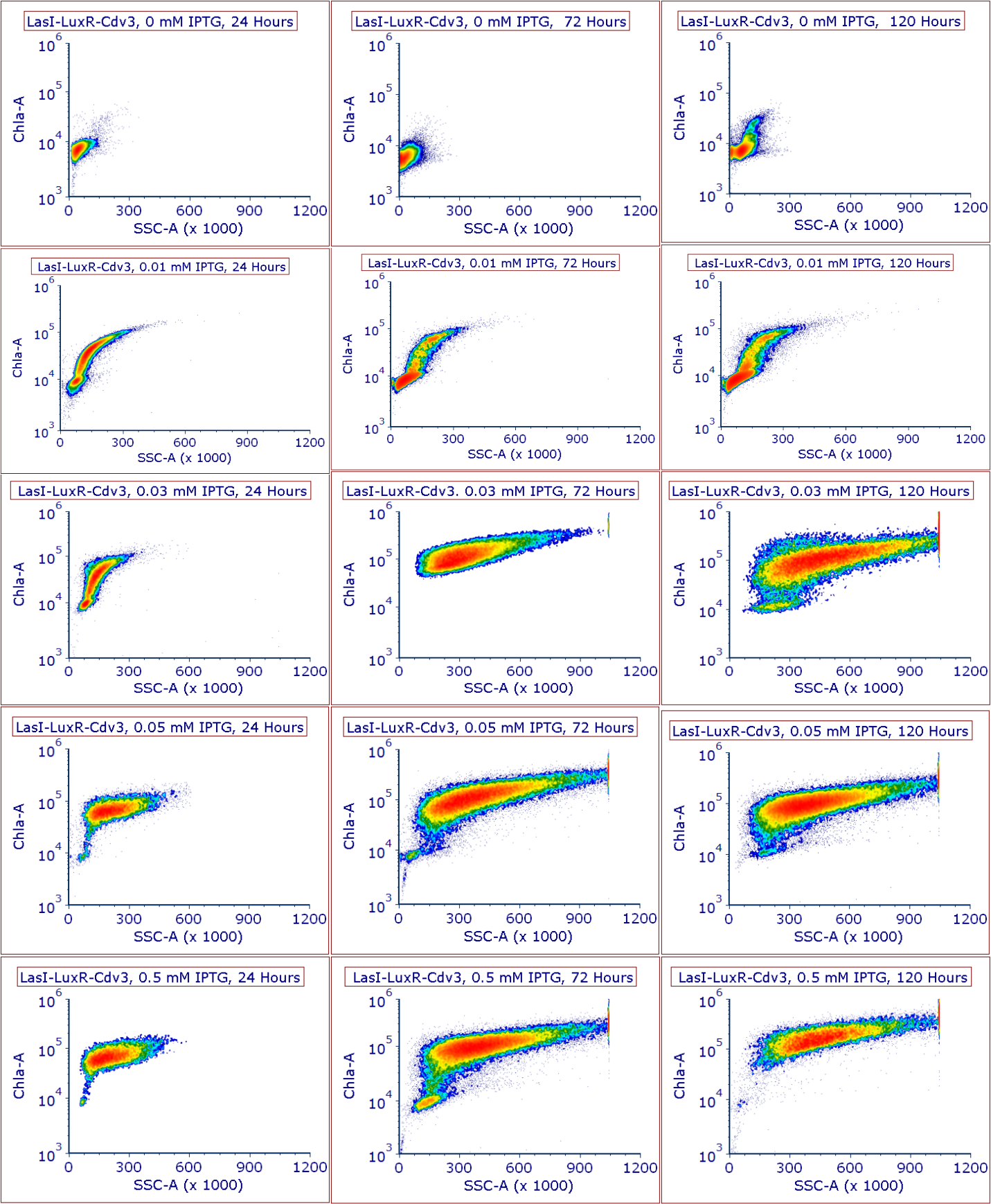

**Supplemental Figure S10.** **Characterization of the LuxR-Cdv3/GFP-LasI strain using flow cytometry.** Cell length monitor of the LuxR-Cdv3/GFP-LasI strain overtime 24,72 and 120 hours, across different IPTG concentrations ranging from 0 up to 0.5 mM of IPTG.Chla-A, chlorophyll *a* area; SSC-A, side scatter area.

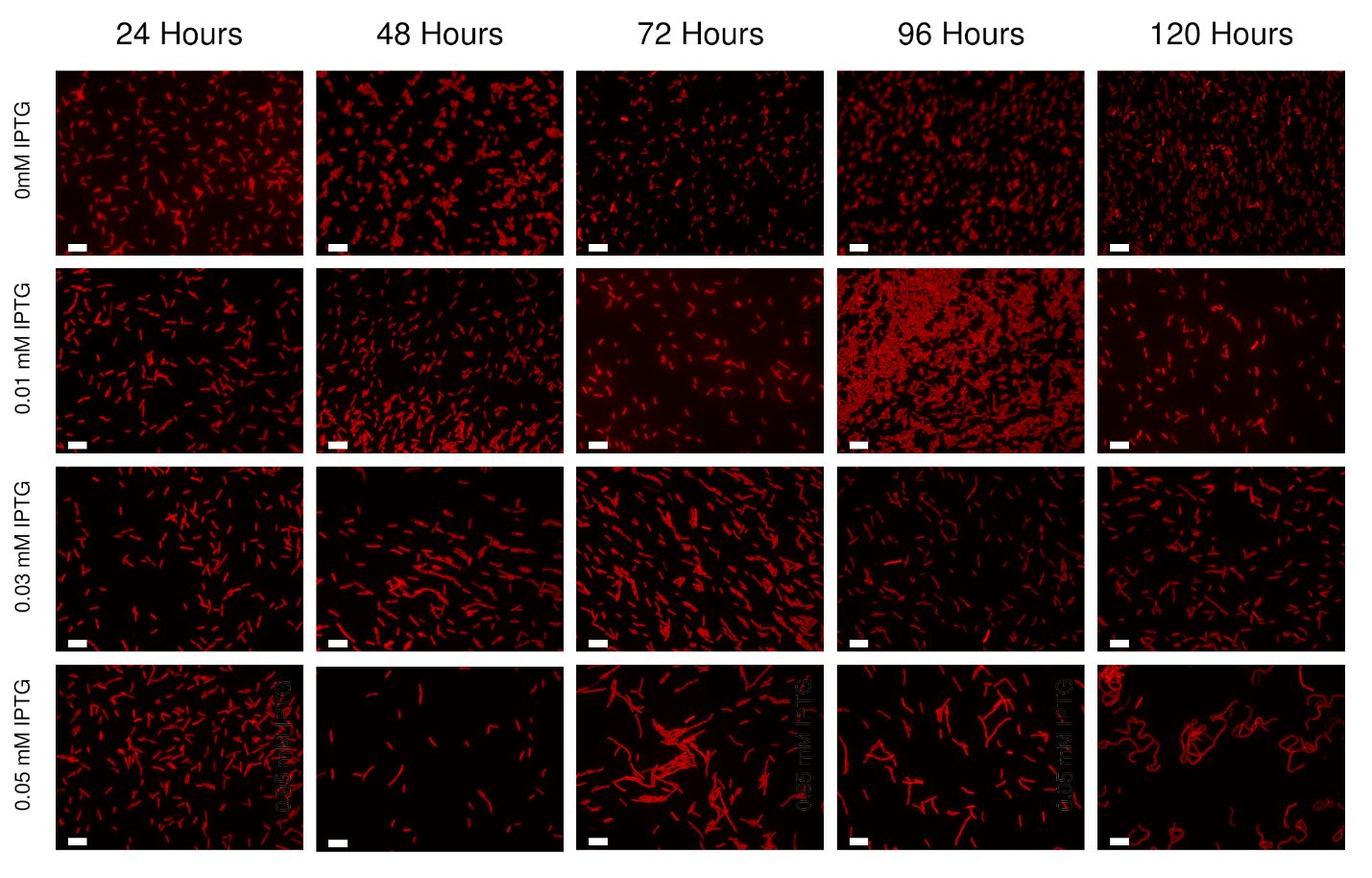

**Supplemental Figure S11: Characterization of the LuxR-Cdv3/GFP-LuxI strain.** Cell length monitor overtime from 24 hours up to 120 hours of the LuxR-Cdv3/GFP-LuxI strain and IPTG concentration ranging from 0 up to 0.05 mM IPTG, using a fluorescence microscope under red channel by tracking the chl *a*. Scale bar: 10 μm.

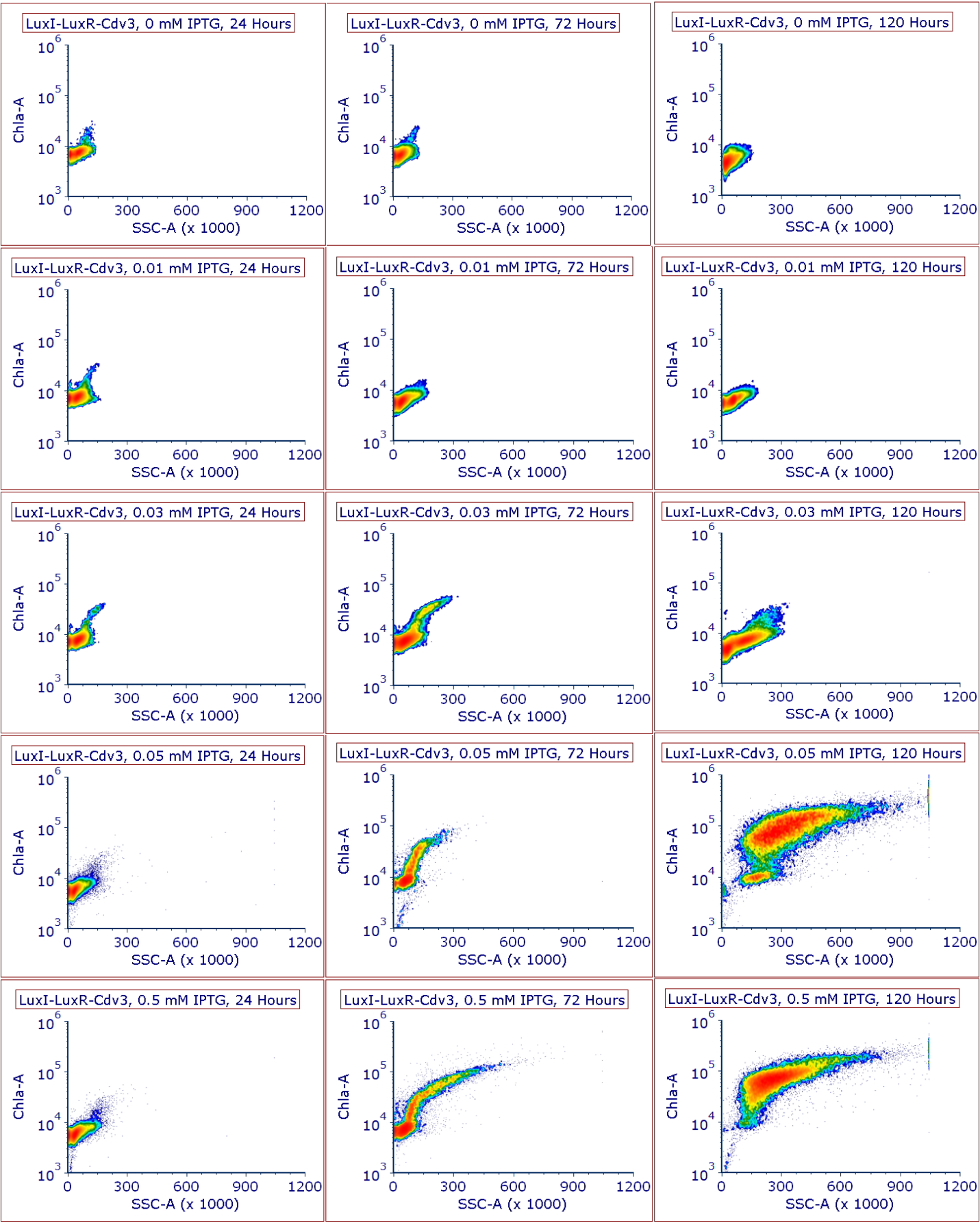

**Supplemental Figure S12. Characterization of the LuxR-Cdv3/GFP-LuxI strain using flow cytometry.** Cell length monitor of the LuxR-Cdv3/GFP-LuxI strain overtime, 24,72 and 120 hours, across different IPTG concentrations ranging from 0 up to 0.5 mM of IPTG. Chla-A, chlorophyll *a* area; SSC-A, side scatter area.

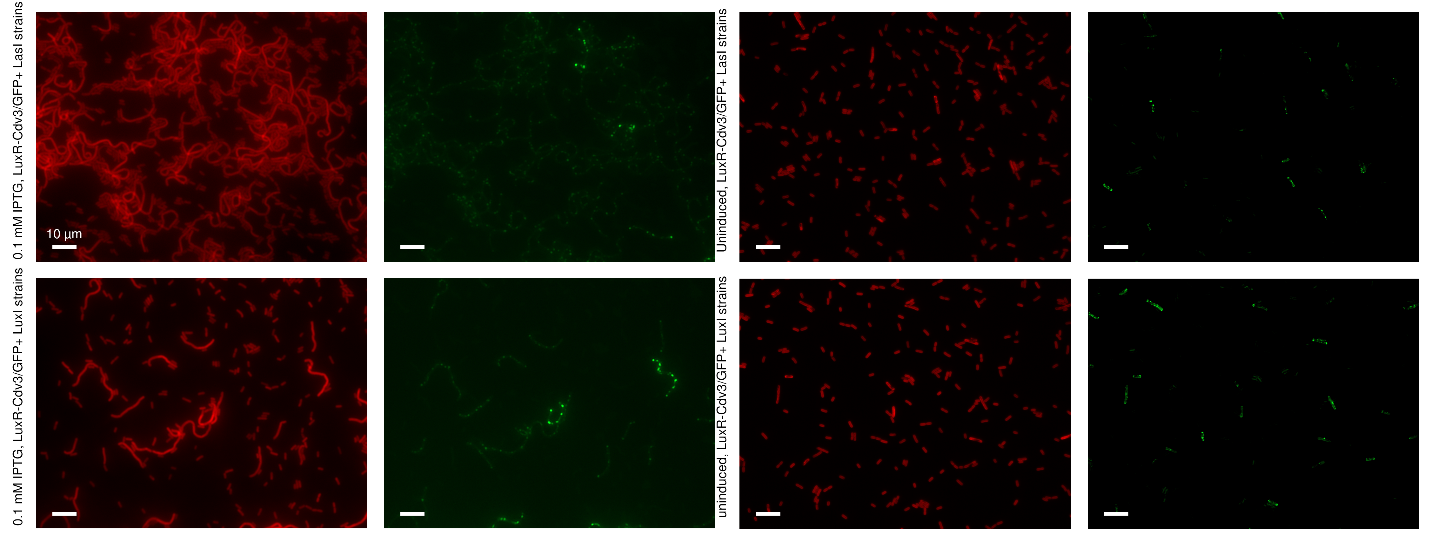

**Supplemental Figure S13. Incubation of sender and receiver strain**. Cell length monitor at 72 hours of the LuxR-Cdv3/GFP+LasI and LuxR-Cdv3/GFP+LuxI strains with 0.1 mM IPTG and uninduced conditions, using a fluorescence microscope under red channel by tracking the chl a and under green channel is the GFP fluorescent reporter. Scale bar: 10 μm.

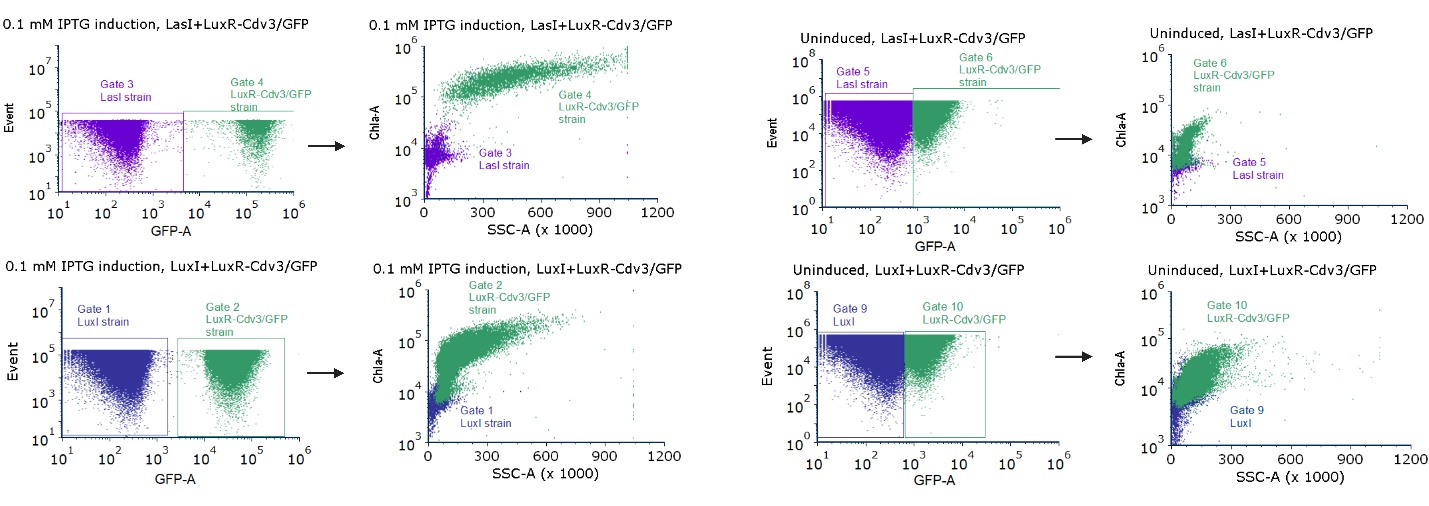

**Supplemental Figure S14. Monitor and differentiation of sender LasI or LuxI with the LuxR-Cdv3/GFP receiver strain using attune flow cytometry.** Cell length monitor at 72 hours of the LuxR-Cdv3/GFP+LasI and LuxR-Cdv3/GFP+LuxI strains with 0.1 mM IPTG and uninduced conditions. With appropriate gating we are able to distinguish the sender and the receiver strain in the same culture, as the receiver strain has a green fluorescent protein. GFP-A, green fluorescent protein area, chlorophyll *a* area; SSC-A, side scatter area.
